# Linking transcriptome and chromatin accessibility in nanoliter droplets for single-cell sequencing

**DOI:** 10.1101/692608

**Authors:** Song Chen, Blue B Lake, Kun Zhang

## Abstract

Linked profiling of transcriptome and chromatin accessibility from single cells can provide unprecedented insights into cellular status. Here we developed a droplet-based Single-Nucleus chromatin Accessibility and mRNA Expression sequencing (SNARE-seq) assay, that we used to profile neonatal and adult mouse cerebral cortices. To demonstrate the strength of single-cell dual-omics profiling, we reconstructed transcriptome and epigenetic landscapes of cell types, uncovered lineage-specific accessible sites, and connected dynamics of promoter accessibility with transcription during neurogenesis.

---

RNA sequencing of single cells or nuclei can reveal their transcription state, while chromatin accessibility sequencing would uncover the associated upstream transcriptional regulatory landscape. Current strategies^1,2^, which involve profiling these modalities separately followed by computational integration, may not fully recapitulate the true biological state. Joint profiling of two layers of -omics information within the same cells would enable a direct matching of transcriptional regulation to its output, allowing for more accurate reconstruction of the molecular processes underlying a cell’s physiology. To enable highly parallel profiling of chromatin accessibility and mRNA from individual nuclei, we developed SNARE-seq, implemented on a micro-droplet platform^3^. In this method, the accessible chromatin in permeabilized nuclei are captured by the Tn5 transposase, prior to droplet generation. We reason that, without heating or detergent treatment, binding of transposases to its DNA substrate after transposition could maintain the contiguity of the original DNA strands^4^, allowing for the co-packaging of accessible genomic sites and mRNA from individual nuclei in the same droplets.

As such, we designed a splint oligonucleotide with sequence complementary to the adapter sequence inserted by transposition (5’ end) and the poly A bases (3’ end) allowing capture by oligo-dT-bearing barcoded beads. After encapsulation of nuclei, their mRNAs and fragmented chromatin can be released by heating the droplets, allowing access to splint oligos and adaptor coated beads having a shared cellular barcode (Fig. 1a). A pair of RNA-seq and chromatin accessibility libraries can then be generated for sequencing. The resulting data can then be connected by their shared cellular barcodes, without the need for probabilistic mapping of single-cell clusters from separate analyses. While SNARE-seq shows similarities to sci-CAR^5^ conceptually, our method was implemented on a widely accessible Drop-seq platform and provides denser chromatin information due to a design that captures chromatin information first, then linking it to the transcriptome.

**Figure 1.**
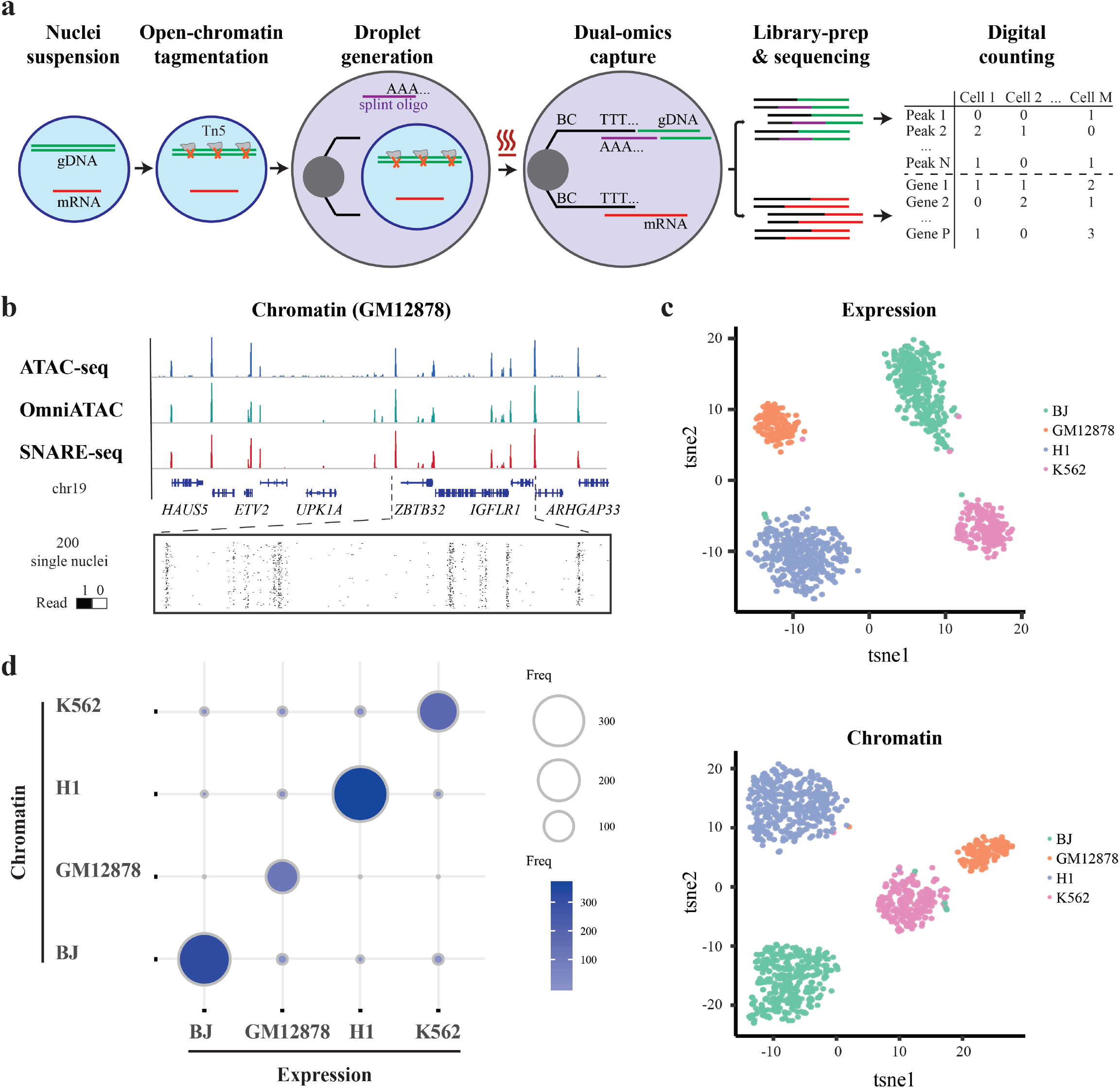
Linked single-nucleus transcriptome and chromatin accessibility sequencing of human cell mixtures. **a**, Workflow of SNARE-seq. Key steps are outlined in the main text. **b**, Aggregate single-nucleus chromatin accessibility profiles recaptured published profiles of ATAC-seq and Omni-ATAC in GM12878 cells. **c**, t-SNE visualization of SNARE-seq gene expression (upper panel) and chromatin accessibility (lower panel) data from BJ, GM12878, H1 and K562 cell mixture. Cellular identities are colored based on independent clustering results with either expression or chromatin data. **d**, Inter-assay identity agreement reveals consistent linked transcriptome and chromatin accessibility profiles of SNARE-seq data. The size and color depth of each circle represents the number of cellular barcodes that were identified by the different assays.

To evaluate SNARE-seq’s ability to capture accessible chromatin, we first performed a proof-of-concept experiment on lymphoblastoid cell line GM12878, which have extensively characterized chromatin landscapes. Ensemble profiles of SNARE-seq accessibility data showed a signal-to-noise ratio similar to ATAC-seq^6^ and Omni-ATAC^7^ (Fig. 1b). The aggregate SNARE-seq data also showed the expected periodical nucleosome pattern and a strong enrichment of fragments within canonical promoter regions (**Fig. S1a, c**), which are typical characteristics of bulk ATAC-seq data. We validated the peaks called from the SNARE-seq data by overlapping them with those of published bulk ATAC-seq and Omni-ATAC data^6^ (**Fig. S1b**) and found that 85.9% of ATAC-seq peaks were shared among all the three assays, and that 87.6% of Omni-ATAC peaks were shared between Omni-ATAC and SNARE-seq. After filtering out low quality data, we obtained a median of 2720 accessible sites per nuclei, which is comparable with several published single cell/nuclei ATAC-seq methods (**Fig. S1d and S2a**).

To assess the accuracy of SNARE-seq in identifying cell types, we performed SNARE-seq on mixtures of cultured human BJ, H1, K562 and GM12878 cells, and collected 1,047 paired profiles (median 500 UMIs; median 805 accessible sites, **Fig. S2a and S3a**). Separate clustering of expression and chromatin accessibility data showed clear separation into four distinct clusters (Fig. 1c). Differential expression of maker genes (**Fig. S4a**) validated these cluster identities. The classification results from both profiles were in good agreement (kappa coefficient of 0.92, Fig. 1d). Notably, we found that transcription factors *JUN, IRF8, POU5F1* and *GATA1*, which showed enriched expression in BJ, GM12878, H1 and K562 cells, respectively (**Fig. S4c**), also exhibited a similar pattern of preferential binding to chromatin sequences captured by SNARE-seq accessibility assay (**Fig. S4d**), consistent with previous observations^8^. We improved the detection sensitivity on chromatin further by using NP40 based Nuclei EZ buffer to boost tagmentation efficiency and adding RNase inhibitor combination^9^ to protect RNA from degradation. From the mixed cell lines we acquired 1,043 paired profiles with median number of 1159 UMIs and 2254 accessible sites captured (**Fig. S2a, S3a**). We then compared SNARE-seq expression and chromatin data with those generated from snDrop-seq or SNARE-seq chromatin only experiments, and observed consistent clustering (**Fig. S5a**,**b**), high level correlation of raw reads (**Fig. S5d**), as well as efficient recovery rate (**Fig. S5e**, 66% recovery of RNA and 100% recovery of chromatin). Furthermore, the species-mixing experiment indicated a high purity and low doublet rate (6%, **Fig. S6**) of SNARE-seq. Therefore, on simple cell mixture, SNARE-seq can effectively separate cell types based on both their chromatin signatures and transcriptomes, with a high level of concordance.

We next applied SNARE-seq to mouse neonatal cerebral cortex (postnatal day 0, n=5) and recovered 5,081 nuclei that had linked transcriptome (median 357 UMIs) and chromatin accessibility (median 2583 accessibility sites) data after QC filtering (**Fig. S2a, S3a**). Correlation analysis of expression or chromatin profiles demonstrated great reproducibility between independent SNARE-seq experiments (**Fig. S7**). Among all RNA reads, 94% aligned to the genome, with 37% of these mapped to exons and 42% mapped to introns (**Fig. S8**), reflecting the enrichment of nascent transcripts in the nucleus^1^. In comparison, despite a similar mapping rate (>91%), the chromatin accessibility data showed a larger fraction of reads (34%) mapped to intergenic regions. There was also enrichment of accessibility reads in close proximity to the transcription start site (10%) and low coverage in exons, suggestive of enhancer and promoter sequences present in those noncoding regions. Therefore, both RNA and chromatin reads showed expected genome distributions comparable to the snDrop-seq^1^ and snATAC data^10^.

Unsupervised clustering of cerebral cortex transcriptomes identified 19 cell clusters, including: astrocytes/radial glia (Ast/RG); intermediate progenitor cells (IP); excitatory neurons (Ex); migrating inhibitory neurons (In); and Cajal-Retzius cells (CR). We further detected several non-neuronal cell types, including: oligodendrocyte progenitor cells (OPC); endothelial cells (End); pericytes (Peri); and microglia (Mic). These cell clusters ranged in size from 37 (0.7%) to 542 (10.7%) cells (**Fig. S9a**), and were independent of batch or sequencing depth (**Fig. S9b-e**). Uniform Manifold Approximation and Projection (UMAP) revealed a trajectory extending from the progenitor states reflective of the sequential development of cortical cell fates. Consistently, nuclei occurring adjacent to intermediate progenitors represented those of the late born neurons of the superficial layers (proceeding deep layer neurons) and glial cell types associated with the onset of gliogenesis that is expected at this time point (Fig. 2a). We compared SNARE-seq transcriptome data with a recently published single-cell RNA-seq dataset of the mouse cortex at a similar developmental time point that was generated by SPLiT-seq^9^. Despite a lower number of detected UMIs, the cell types and their signatures were reasonably well correlated (**Fig. S10a-c**). Notably, we also captured finer distinctions between closely related cellular states and identified three sub-clusters of intermediate progenitor cells: cluster IP-Hmgn2, expressing *Mki67*, *Top2a* and *Kif23* (Fig. 2b, **Fig. S10b** and **Table S1**), representing cycling progenitors; cluster IP-Gadd45g, which was enriched for *Gadd45g*, representing apical progenitors that exited from cell-cycle^11^; and cluster IP-Eomes, representing basal progenitors that show early commitment to the neuronal lineage. Cell-type and layer identities of our clusters were further validated by expected expression of known marker genes and *in situ* staining of novel discovered makers (Fig. 2b, **S11 and Table S1**).

**Figure 2.**
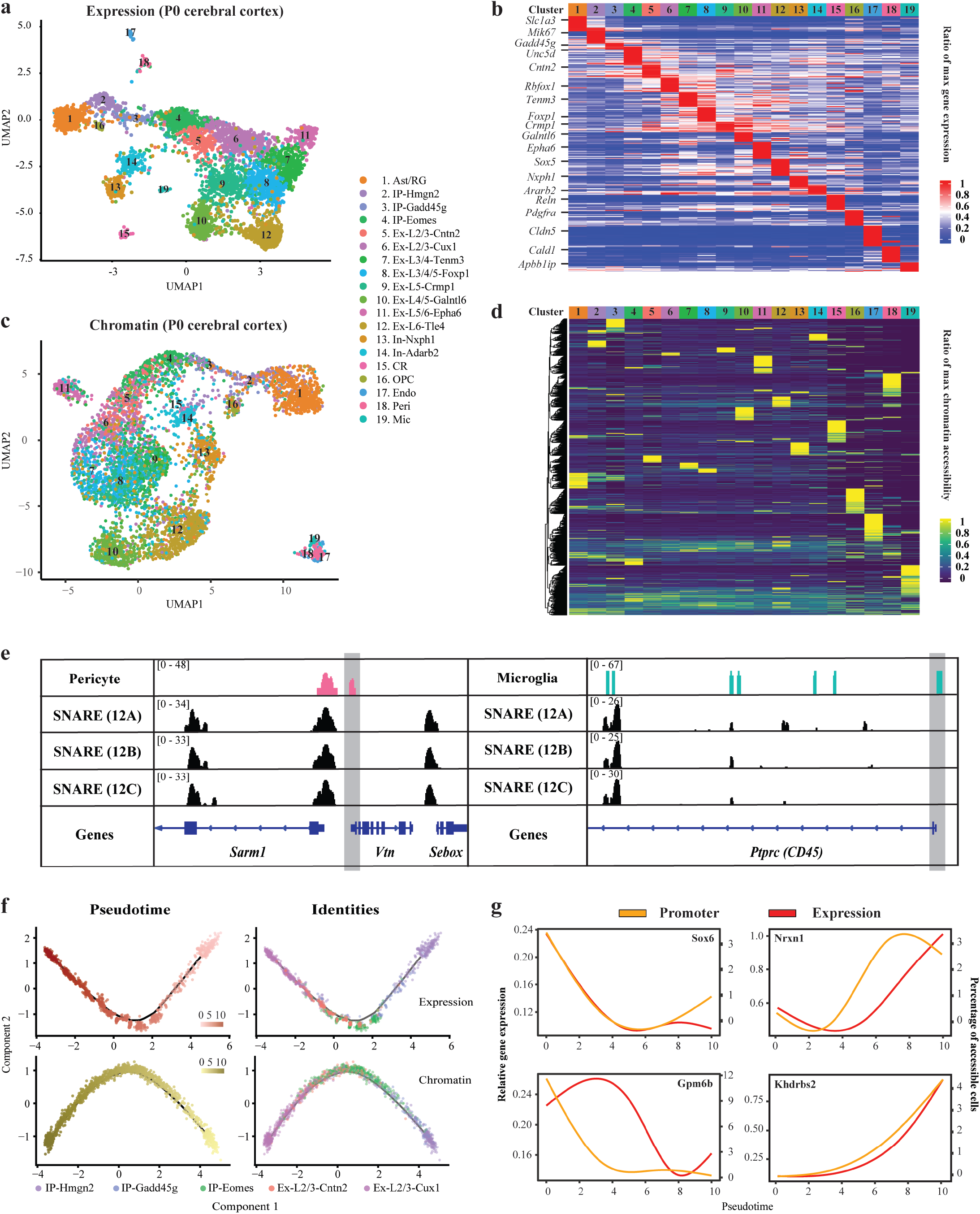
Dual-omics profiling of neonatal mouse cerebral cortex with SNARE-seq. **a**, UMAP projection of SNARE-seq expression data of mouse cerebral cortex nuclei. Cell types were assigned based on known markers. **b**, Heatmap showing the normalized expression of cell type-specific genes relative to the maximum expression level across all cell types. **c**, UMAP projection of SNARE-seq chromatin accessibility data of mouse cerebral cortex nuclei. Cells are labeled with the same color codes for cell types identified by the linked expression data. **d**, Heatmap showing the normalized chromatin accessibility of type-specific accessible sites relative to the maximum accessibility across all cell types. **e**, Chromatin accessibility tracks generated from cell-type specific or batch aggregated (batch code 12A, 12B and 12C) chromatin accessibility data at pericyte (left) and microglia (right) marker gene loci (*Vtn* and *CD45* respectively). For better visualization, the promoter regions were shaded in gray. **f**, Pseudotime trajectories constructed with SNARE-seq expression (upper panels) and chromatin accessibility (lower panels) profiles for 1,469 nuclei (214 IP-Hmgn2, 99 IP-Gadd45g, 437 IP-Eomes, 177 Ex-L2/3-Cntn2 and 542 Ex-L2/3-Cux1) from the mouse cerebral cortex. Cells are colored according to pseudotime score (left panels) or cellular identities (right panels). **g**, Promoter accessibility (yellow) and gene expression (red) changes of *Sox6*, *Gpm6b*, *Nrxn1* and *Khdrbs2* across pseudotime during early neurogenesis. **Misc**, cells of miscellaneous clusters.

We compared aggregated SNARE-seq chromatin accessibility profiles with published bulk ATAC-seq ENCODE data on neonatal mouse brain cortex and found a strong concordance between these two methods (**Fig. S10e-f**). To cluster co-assayed cells based on their chromatin accessibility profiles, we used their corresponding transcriptional profiles to aggregate chromatin accessibility signals for each cluster separately, followed by peak calling and the probabilistic topic modeling method implemented in cisTopic^12^. After projecting onto lower dimensions using UMAP, most single-nuclei chromatin accessibility clusters (Fig. 2c), corresponded to the same cell types resolved from the corresponding expression data (Fig. 2a). Notably, the chromatin accessibility of deep layer excitatory neurons and migrating inhibitory neurons, which differentiated earlier in the cerebral cortex and ganglionic eminences, respectively, showed well-separated clusters, whereas those of late-generated superficial layer excitatory neurons were less distinct. Interestingly, those diffuse boundaries identified by expression profile were also clustered as subtypes based on chromatin information (**Fig. S12a-b**). Those subtypes may represent datasets with insufficient clustering power due to the sparsity of chromatin data and/or dynamic epigenetic states that are still undergoing maturation. Cell-type identities of the major clusters were further supported by the specific accessibility in the promoter region for marker gene loci *Hes5* (Ast/RG), *Gadd45g* (IP), *Meg3* (Neurons), *Pdgfra* (OPC), *Vtn* (Peri) and *Apbb1ip* (Mic) (**Fig. S13**). Importantly, we found that the promoter accessibility of lineage markers *Vtn* and *CD45* (for pericyte and microglia representing 1% and 0.7% of total cells) were present only in cell-type aggregated chromatin profiles that were identified *de novo* with transcriptome data. In contrast, chromatin data analyzed based on the accessible peaks called from the batch-aggregated profiles, the current default strategy for analyzing sc-ATAC-seq data, failed to recover these accessible peaks unique to rare cells in the presence of background noise from other more abundant cells (Fig. 2e). Consistent with this notion, clustering of chromatin profiles generated without using any expression information yielded less clear cell type boundary and many of those low-abundant cell types were largely undetected (**Fig. S12c**). Therefore, *a priori* knowledge of cell type identity in chromatin accessibility data using the linked gene expression profiles permits more sensitive detection of accessible chromatin region. This underscores the strength of our SNARE-seq dual-omic assay over independent single-cell RNA and chromatin accessibility sequencing methods for detecting cell-type and subtype specific gene expression and accessible chromatin.

Differential accessibility (DA) test of SNARE-seq chromatin profiles identified 35,166 sites (p<0.05) across the 19 murine cerebral cortex cell types (Fig. 2d and **Table S2**). Of all 35,166 differential accessible sites, 2,835 (8%) located within promoter regions, and 128 also showed differential gene expression between clusters (**Fig. S14**). For theses 128 genes, the expression levels and their promoter accessibilities across all cell types were mostly positively correlated (median r 0.34, **Fig. S15**), indicating a direct linkage of chromatin accessibilities to the corresponding transcriptomes. To further characterize the DA sites, we performed gene ontology enrichment and motif discovery analysis using GREAT and HOMER, respectively (**Fig. S16**). Notably, genomic elements that were mostly associated with Ast/RG and OPC cells fell into the biological processes regulating stem cell maintenance and differentiation. These sites were further enriched for binding motifs of LHX2 and SOX2, both of which are known regulators of neurogenesis and gliogenesis^13,14^. We also found that differential accessible sites of the IP-Gadd45g cells (representing 1.9% of the total cells) were enriched for the Wnt signaling pathway components, consistent with the role or this pathway in regulating cell cycle exit and promoting neuronal differentiation of intermediate progenitors^15^. Therefore, linking chromatin accessibility profiles to transcriptomic data directly allowed us to effectively identify cell-type specific transcription regulatory mechanisms.

To further demonstrate the utility of having a direct linkage between transcriptome and chromatin accessibility, we focused on the transition of intermediate progenitors to upper layer excitatory neurons. Using Monocle, we ordered gene-expression profiles of 1,469 nuclei along a pseudotime trajectory based on the top differential expressed genes (qval<0.05, Fig. 2f, upper panel). From transcription kinetics, we found a clear pattern originating from a cell-cycle exited state (*Mki67* and *Gadd45g*), that progressed from neuroblast stages (*Eomes* and *Unc5d*) to *Foxp1* and *Cux1*-expressing upper layer neurons^16,17^ (**Fig. S17a**). We further oriented accessibility profiles of the same nuclei along a separate trajectory (Fig. 2f, lower panel) based on a set of 1,332 sites that showed differential accessibility (qval<0.1). Notably, these separately constructed developmental trajectories showed high correlation (r=0.87) along pseudotime. From these differential accessible sites, 103 were found within promoter regions and 21 associated genes that were also differentially expressed by pseudotime. Intriguingly, most of these genes presented similar directional changes in promoter accessibility and expression level (**Fig. S17b-c**). For example, *Sox6*, a transcription factor required for maintenance of neural precursor cells^18^, and membrane protein-encoding *Mlc1* showed a decline along neuronal differentiation, while *Khdrbs2* (SLM1), encoding an RNA-binding protein participating in alternative splicing, and its regulating target encoded by *Nrxn1*^19^ showed similar directional raise along neurogenesis (Fig. 2g, **S18a-b**). Thus, SNARE-seq provided linked expression and chromatin accessibility profiles that enabled construction of regulatory dynamics during developmental programs, as well as detailed characterization of the epigenetic states for these cell clusters (**Fig. S19**).

To demonstrate the effectiveness for SNARE-seq to resolve the more matured and distinct cell states, we additionally processed adult mouse cerebral cortex to obtain 10,309 paired profiles (median 1,332 RNA UMIs and median 2,000 chromatin accessibility sites per nucleus) after QC filtering (**Fig. S2a, S3a**). Unsupervised clustering of the 10,309 transcriptomes revealed 22 cell clusters, including 10 excitatory neuron types, 4 inhibitory neuron types (*Pvalb*, *Sst*, *Npy* and *Vip*-expressing) and oligodendrocyte progenitor cells (OPC), newly-formed *Itpr2*-expressing oligodendrocyte (Oli-Itpr2) and mature oligodendrocyte (Oli-Mal), as well as other non-neuronal cells (Fig. 3a). Most of the clusters can be identified using existing lineage or cortical layer markers, but in a more restricted pattern (**Fig. S11, 20, 21a-c and Table S1**) compared to the neonatal clusters (**Fig.**). To investigate the epigenetic patterns of each cell cluster, we aggregated SNARE-seq chromatin data of the adult cerebral cortex, which showed high similarity to bulk ATAC-seq data (**Fig. S21d**), based on cell-type identifies defined by *de novo* clustering of linked transcriptome data. We then performed peak-calling and clustering using the topic modeling method^12^. The cell clusters were more cleanly and distinctly separated, compared to the chromatin profiles of neonatal cortex, likely due to the more discrete cell states of the adult brain. We next performed gene ontology and motif enrichment analysis on those differential accessible peaks that were identified across all cell clusters (**Table S3**). Although some clusters, such as for the astrocytes and microglia, showed similar enrichment of biological processes and transcription factors (**Fig. S22**), most other clusters revealed regulation of features that were distinct from the corresponding cells in the developing mouse cortex, reflective of their postnatal maturation states within the brain.

**Figure 3.**
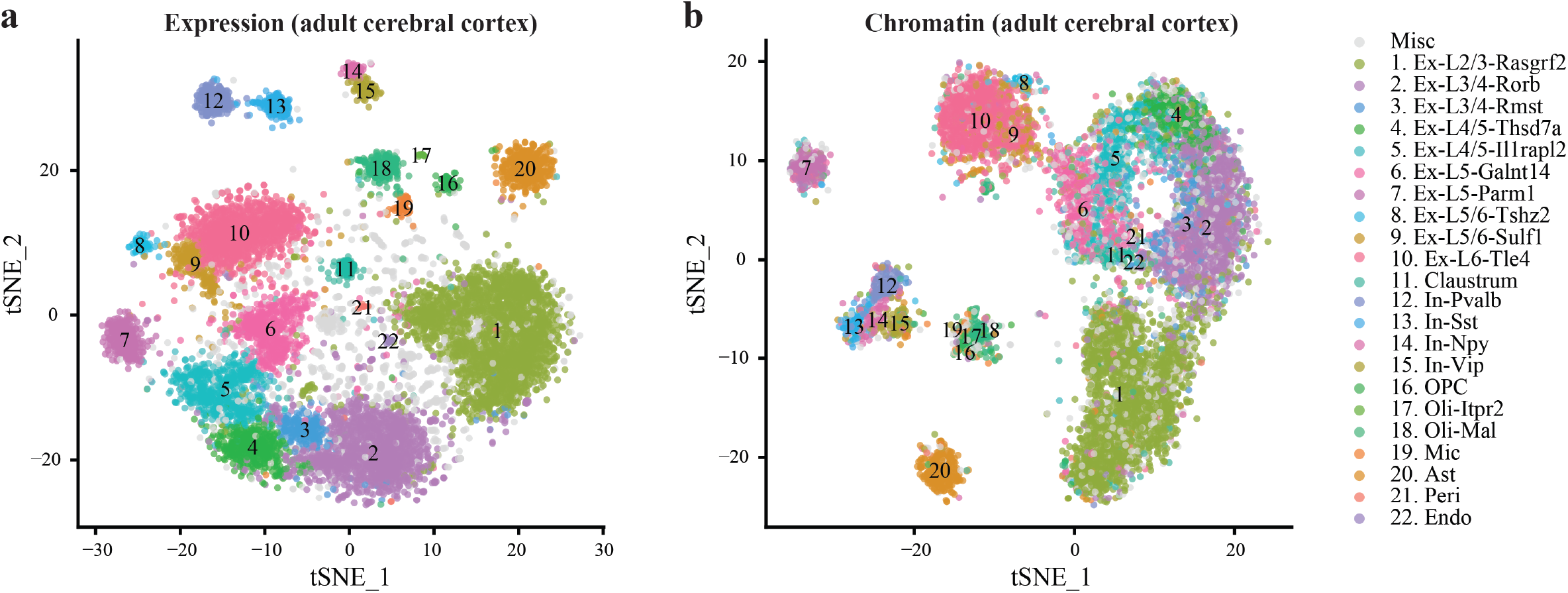
SNARE-seq profiling of adult mouse cerebral cortex. **a**, tSNE projection of SNARE-seq expression data of mouse cerebral cortex 10,309 nuclei. Cell types were assigned based on known markers. **b**, tSNE projection of SNARE-seq chromatin accessibility data of adult mouse cerebral cortex nuclei. Cells were labeled with the same color codes for cell types identified by the linked expression data.

Overall, SNARE-seq is a robust method allowing the joint measurement of transcriptome and chromatin accessibility in single cells or nuclei. Due to a simple design that does not rely on proprietary reagents, SNARE-seq can be widely implemented. Compared to the recently reported sci-CAR^5^, SNARE-seq detects RNA molecules at a sensitivity comparable to other single nucleus RNA-seq methods (**Fig. S3a-b**), captures more accessible sites (∼4-5 times, **Fig. S2a-b**) which allows improved discovery of differentially accessible sites (∼2-fold) and better resolution of cell clusters (**Fig. S19**). Finally, the throughput of this assay could potentially be further improved through integration with a cellular combinatorial indexing strategy^10^. SNARE-seq represents a valuable tool for characterizing tissue complexity through profiling both the inputs and outcomes of transcriptional regulatory units, which would be especially valuable for creating molecular cell atlases of human tissues and clinical samples.

## Acknowledgments

This project was supported by NIH grants U01MH098977, R01HL123755, U54HL145608 to K.Z.

