## Supplementary figures for "Linking transcriptome and chromatin accessibility in nanoliter droplets for single-cell sequencing"

#### Supplementary Figure 1

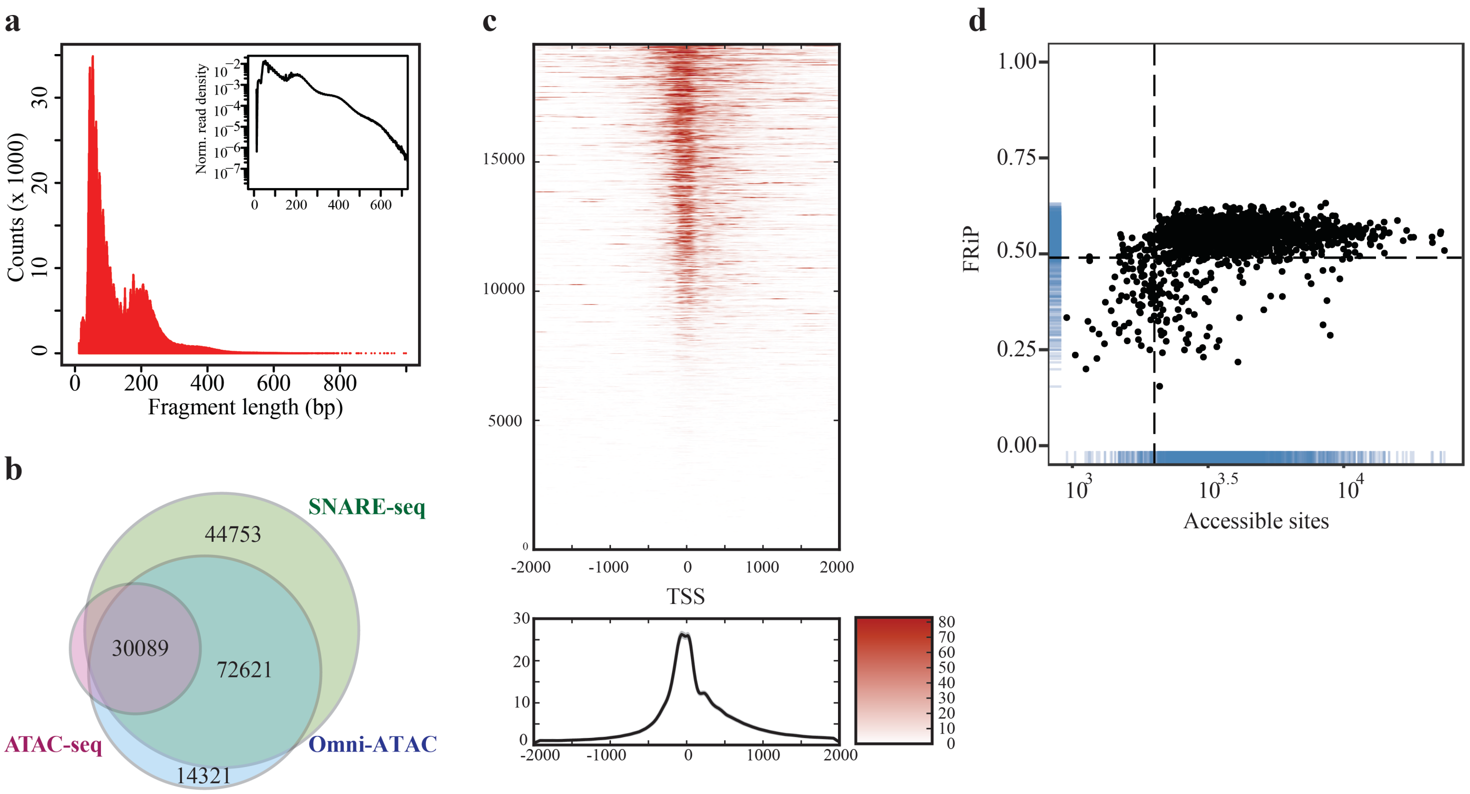


**Supplementary Figure 1.** Quality metrics of SNARE-seq chromatin profiles of GM12878. **a**, DNA fragment size distribution showing nucleosomal periodicity. (Inset) log-transformed histogram. **b**, Representative Venn diagram showing the numbers of overlapping accessible sites generated using ATAC-seq, Omni-ATAC and SNARE-seq chromatin assays. **c**, Enrichment of chromatin accessibility signals around transcription start sites (TSS). Top: enrichment around individual TSS. Bottom: aggregated enrichment across all TSSs. X axis indicates the relative distance to TSSs. **d**, Total number of accessible sites versus fraction of reads in open chromatin peaks (FRiP) within GM12878 cells (n = 3,787 after filtration). Dotted lines (2,000 and 49%) represent cutoffs used for downstream analysis.

### Supplementary Figure 2

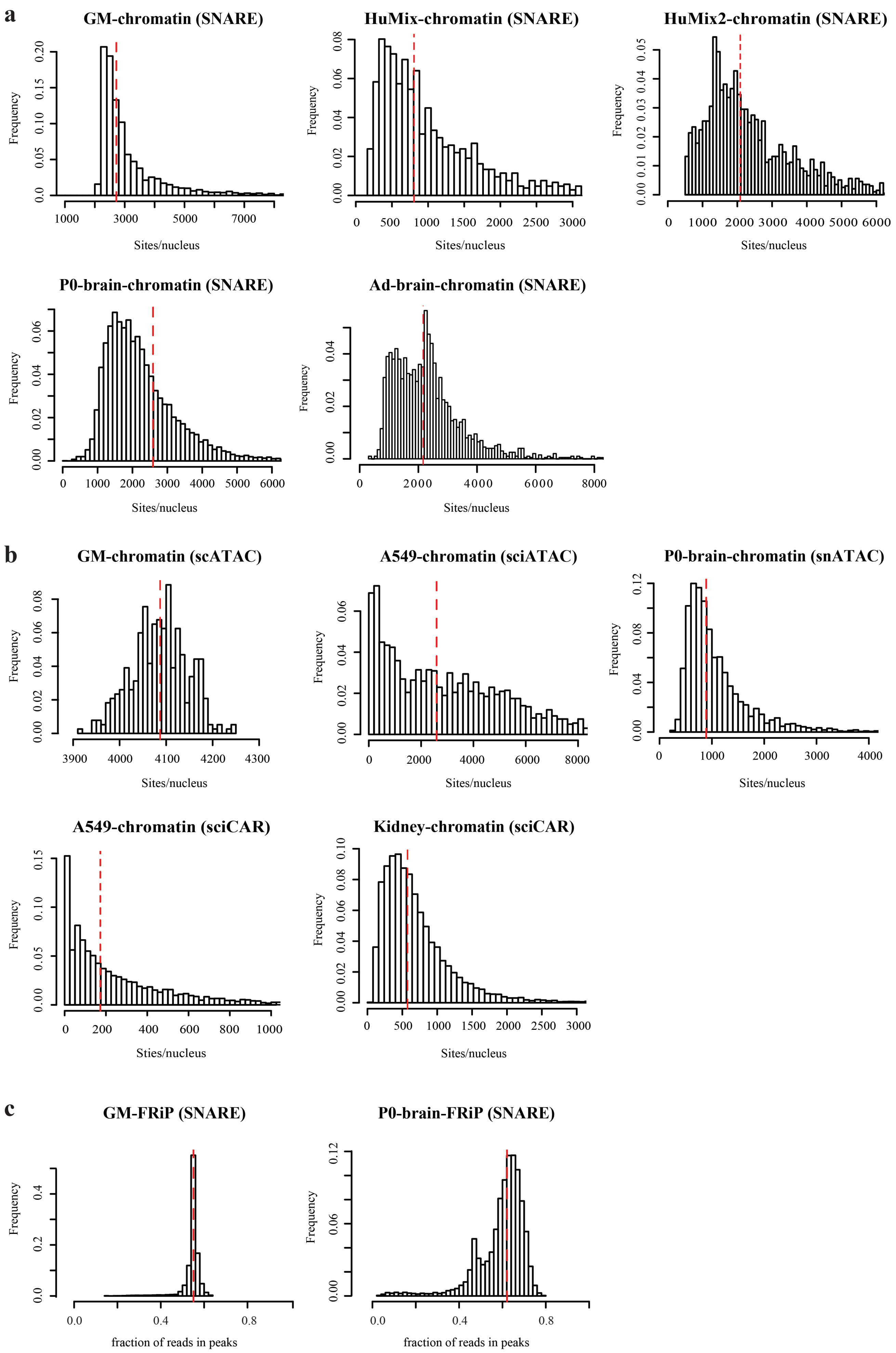

**Supplementary Figure 2.** Comparison of the number of accessible sites detected per nucleus with different single-cell/nucleus chromatin accessibility methods. **a**, Histogram showing the numbers of accessible sites captured by SNARE-seq chromatin profiles. **b**, Histogram showing the numbers of accessible sites detected per nucleus with different single-cell/nucleus ATAC-seq methods. The processed peak count matrices of published reports were downloaded from GEO (scATAC, GSE65360; sci-ATAC, GSE68103; snATAC, GSE100033; sci-CAR, GSE117089) and binarized. **c**, Histogram showing the fraction of reads in peaks (FRiP) within GM12878 or postnatal day 0 mouse cerebral cortex SNARE-seq chromatin accessibility data. GM12878, **GM**; Human cell lines mixture (BJ, GM12878, H1 and K562), lyzed by Triton-X, **HuMix**; Human cell lines mixture, lyzed by Nuclei EZ Prep, **HuMix2**; Postnatal day 0 mouse cerebral cortex, **P0-brain**; Adult mouse cerebral cortex, **Ad-brain**.

### Supplementary Figure 3

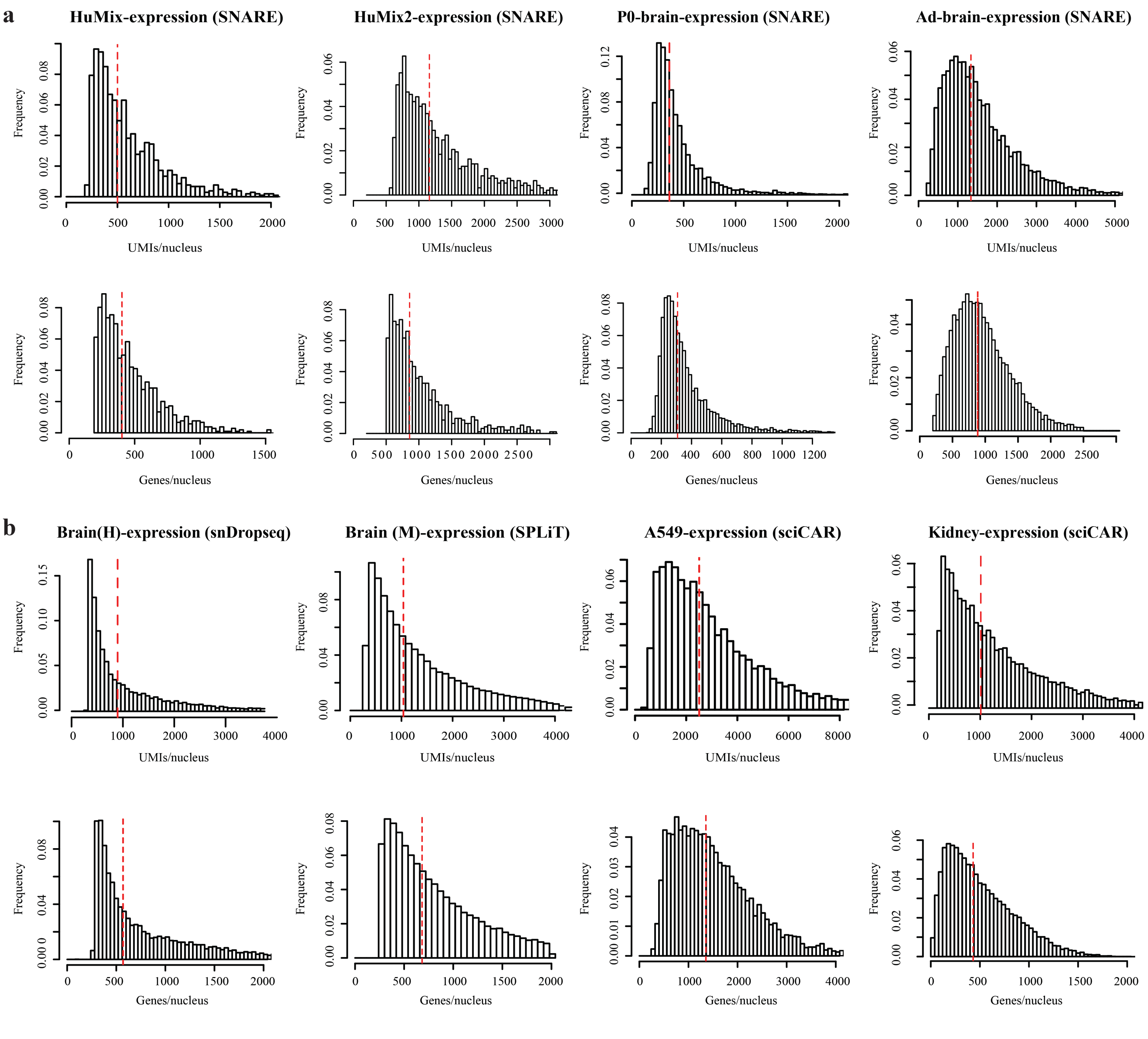


**Supplementary Figure 3.** Comparison of the number of genes and transcripts detected per nucleus with different single-cell/nucleus RNA-seq methods. **a**, Histogram showing the numbers of UMIs and genes captured by SNARE-seq expression profiles. **b**, Histogram showing the number of UMIs and genes detected per nucleus with different single-cell/nucleus RNA-seq methods. The UMI count matrices of published reports were downloaded from GEO (snDrop, GSE97942; SPLiT-seq, GSE110823; sciCAR, GSE117089). Adult human brain cortex, **Brain (H)**; Postnatal day 2 mouse cerebral cortex, **Brain (M)**.

#### Supplementary Figure 4

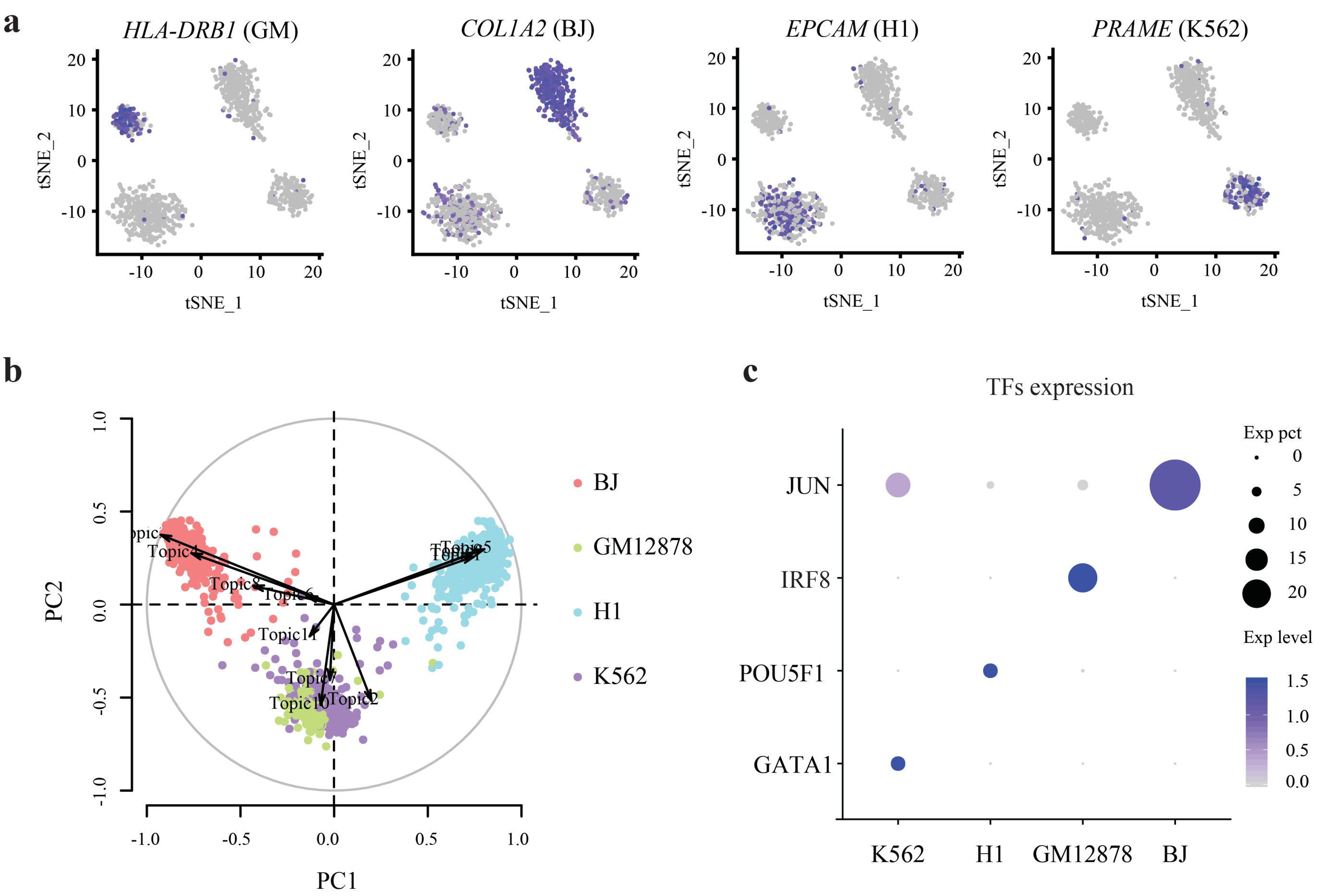


**d**

| TFs \ Cells | Motif | K562 | H1 | GM12878 | BJ |
|---|---|---|---|---|---|
| JUN | 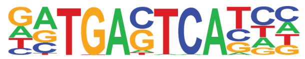 | 1.00E-10 | N.D. | 1.00E-61 | 1.00E-140 |
| IRF8 | 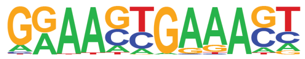 | 1.00E-02 | N.D. | 1.00E-30 | 1.00E-02 |
| POU5F1 | 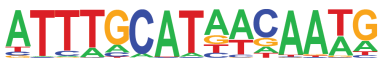 | N.D. | 1.00E-26 | 1.00E-02 | N.D. |
| GATA1 | 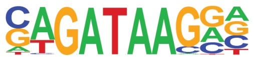 | 1.00E-06 | N.D. | N.D. | N.D. |

**Supplementary Figure 4.** SNARE-seq identified cell types within human cell line mixture. **a**, Feature plot showing the marker gene expression of individual cell lines within each cluster. **b**, Biplot showing the contribution of accessible peak topics identified by cisTopic in classifying cell types with chromatin data. **c**, Dot plot showing the expression of transcription factors (TF) in individual clusters. The size of the dot represents the percentage of nuclei within a cell type expressing the transcription factor and the color indicates the average expression level. **d**, Motif analysis identified the level of significance (in p-value) of transcription factor binding within differential accessible peak topics as mentioned above. p-value of marker TF for each cell type is colored in red.

#### Supplementary Figure 5

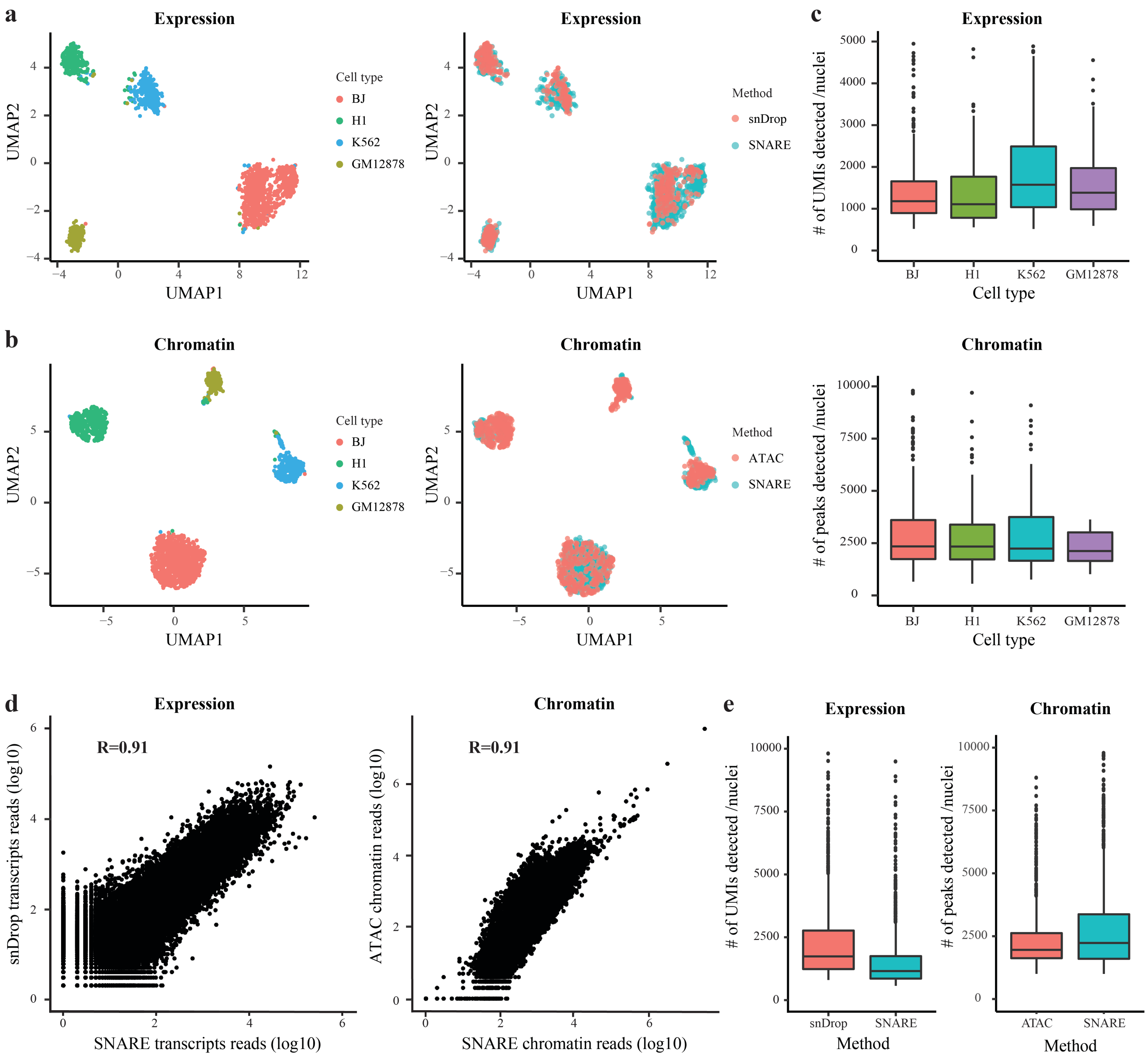


**Supplementary Figure 5.** Comparison of SNARE-seq dual-omics assay with single-omic expression (snDrop-seq) and chromatin (chromatin only) methods. **a**, Clustering of snDrop-seq and SNARE-seq combined expression profiles of human cell line mixture. Cells were labeled by cell type (left) or method (right). **b**, Clustering of SNARE-seq chromatin profiles (dual or chromatin-only assay) of human cell line mixture. Cells were labeled by cell type (left) or method (right). **c**, Distribution of transcripts and accessible chromatin peaks detected by SNARE-seq method in individual cell types. **d**, Correlation of gene expression and chromatin profiles between dual- and single-omic assays. Aggregated transcript reads and chromatin reads were log10 normalized. **e**, Distribution of transcripts and chromatin peaks detected by dual- and single-omic assays. The median numbers of transcripts detected by snDrop-seq and SNARE-seq are 1747 and 1159 respectively and the median number of chromatin peaks detected by SNARE-seq single- and dual-omic assay are 2254 and 1960 respectively.

#### Supplementary Figure 6

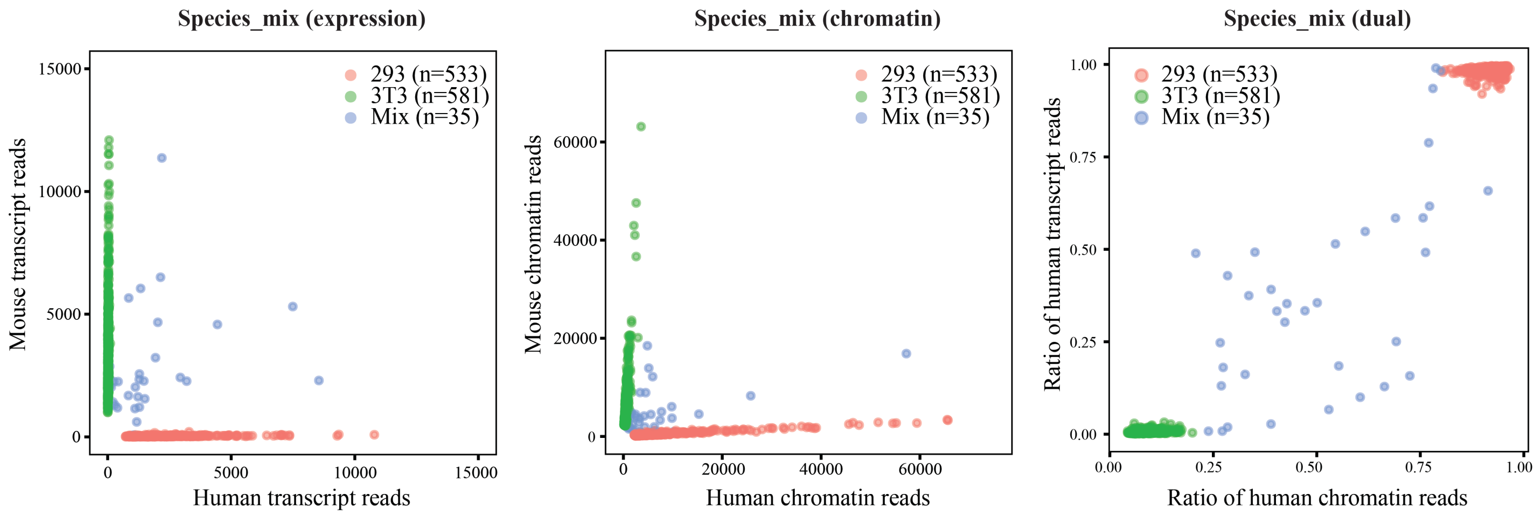


**Supplementary Figure 6.** Species-mixing experiment showing the transcript and chromatin reads detected by SNARE-seq and proportion of human reads in each barcodes.

**Supplemetnary Figure 7**

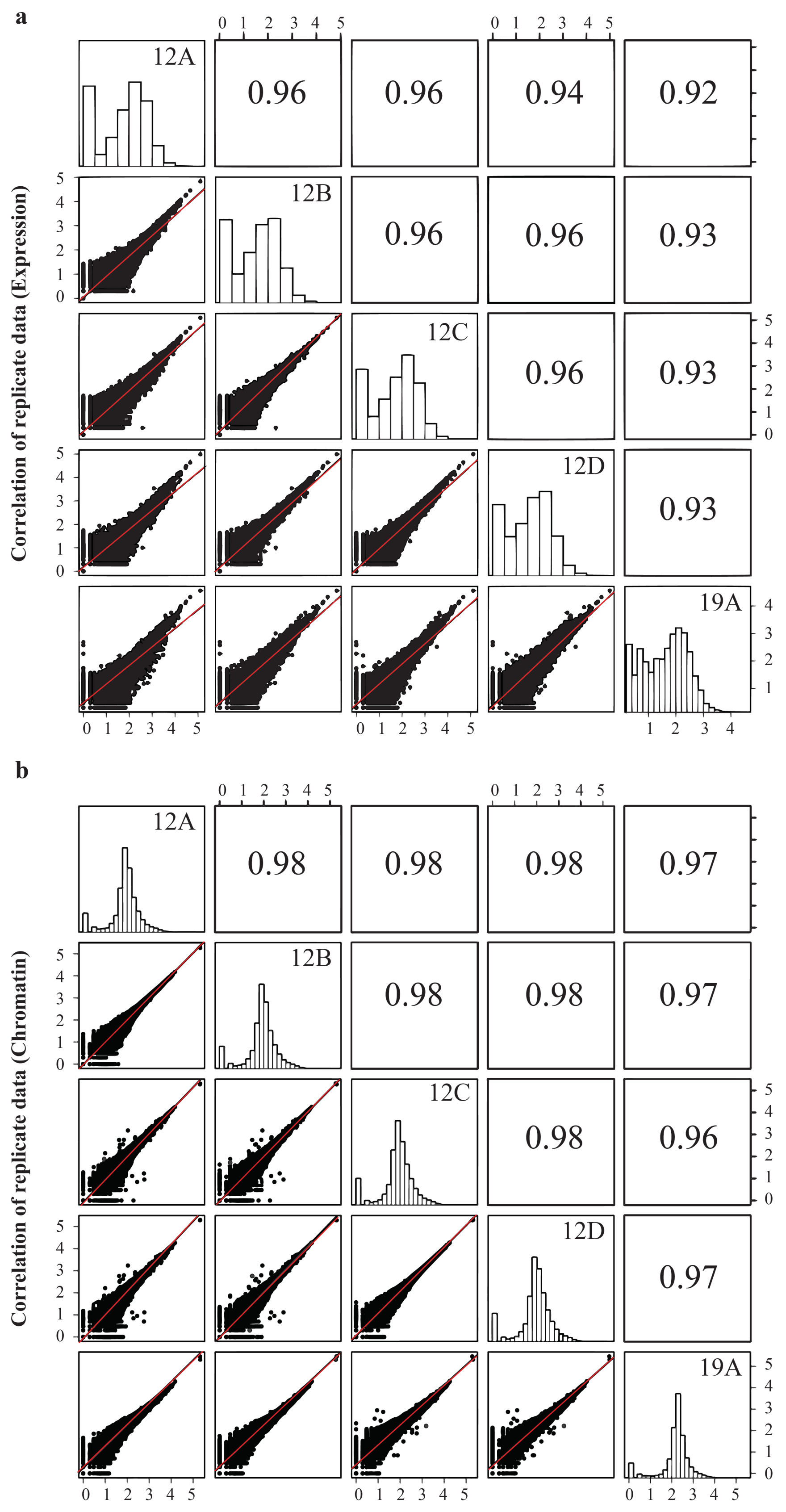


**Supplementary Figure 7.** Reproducibility of SNARE-seq. **a**, Pair-wise correlation of gene expression profiles between individual replicates of postnatal day 0 sample. Aggregated transcript reads were log10 normalized. **b**, Pair-wise correlation of chromatin accessibility profiles between individual replicates. Aggregated genome coverage was log10 normalized.

#### Supplementary Figure 8

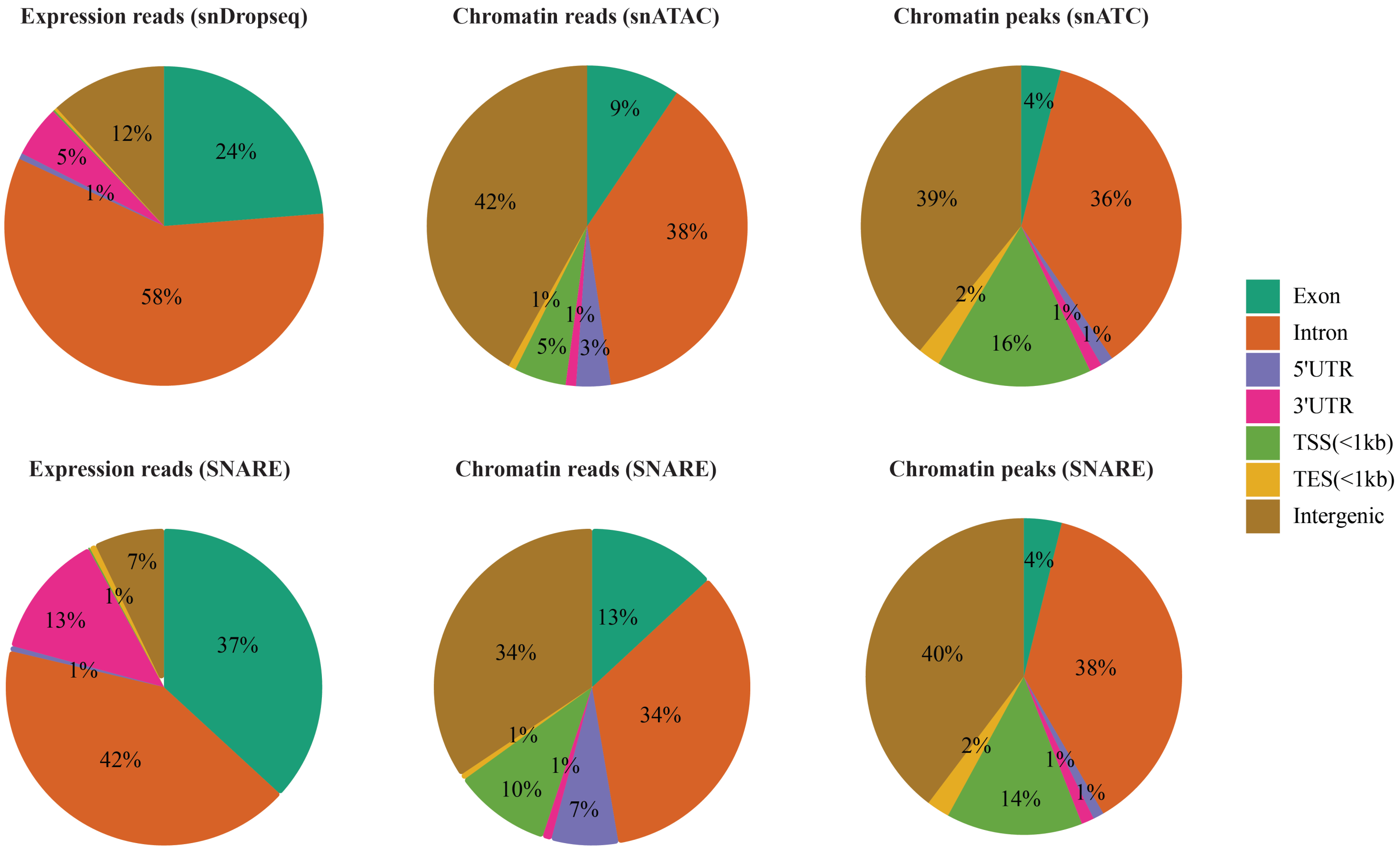


**Supplementary Figure 8.** Proportion of sequencing reads mapped to different genomic features. Top, mapping of reference expression reads[2], chromatin reads[17] and accessible peaks. Bottom, mapping of SNARE-seq expression reads, chromatin reads and accessible peaks of mouse cerebral cortex data. For this analysis, total expression reads of snDrop-seq and SNARE-seq are 32,059,445 and 8,238,261, respectively. Total chromatin reads and peaks called are 180,548,727 and 140,102, 428,942,515 and 175,298 for snATAC and SNARE-seq, respectively.

**Supplementary Figure 9**

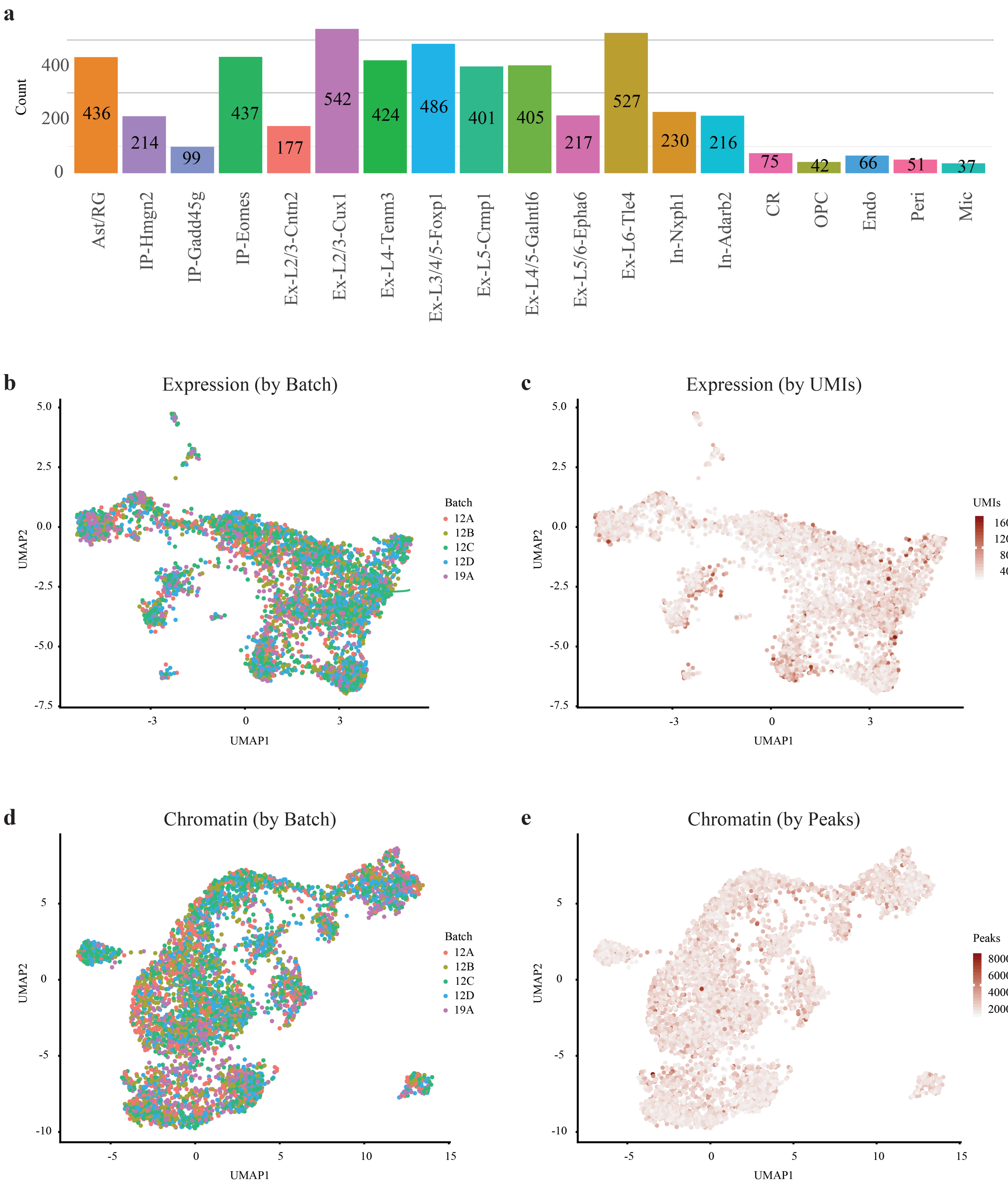


**Supplementary Figure 9.** Robustness of SNARE-seq. **a**, Barplot showing the numbers of nuclei recovered for each cell type. UMAP projection of mouse cerebral cortex expression data as in Fig. 2a showing batch identity (**b**), and UMI read depth (**c**). UMAP projection of chromatin accessibility data as in Fig. 2c showing batch identity (**d**), and peak read depth (**e**).

#### Supplementary Figure 10

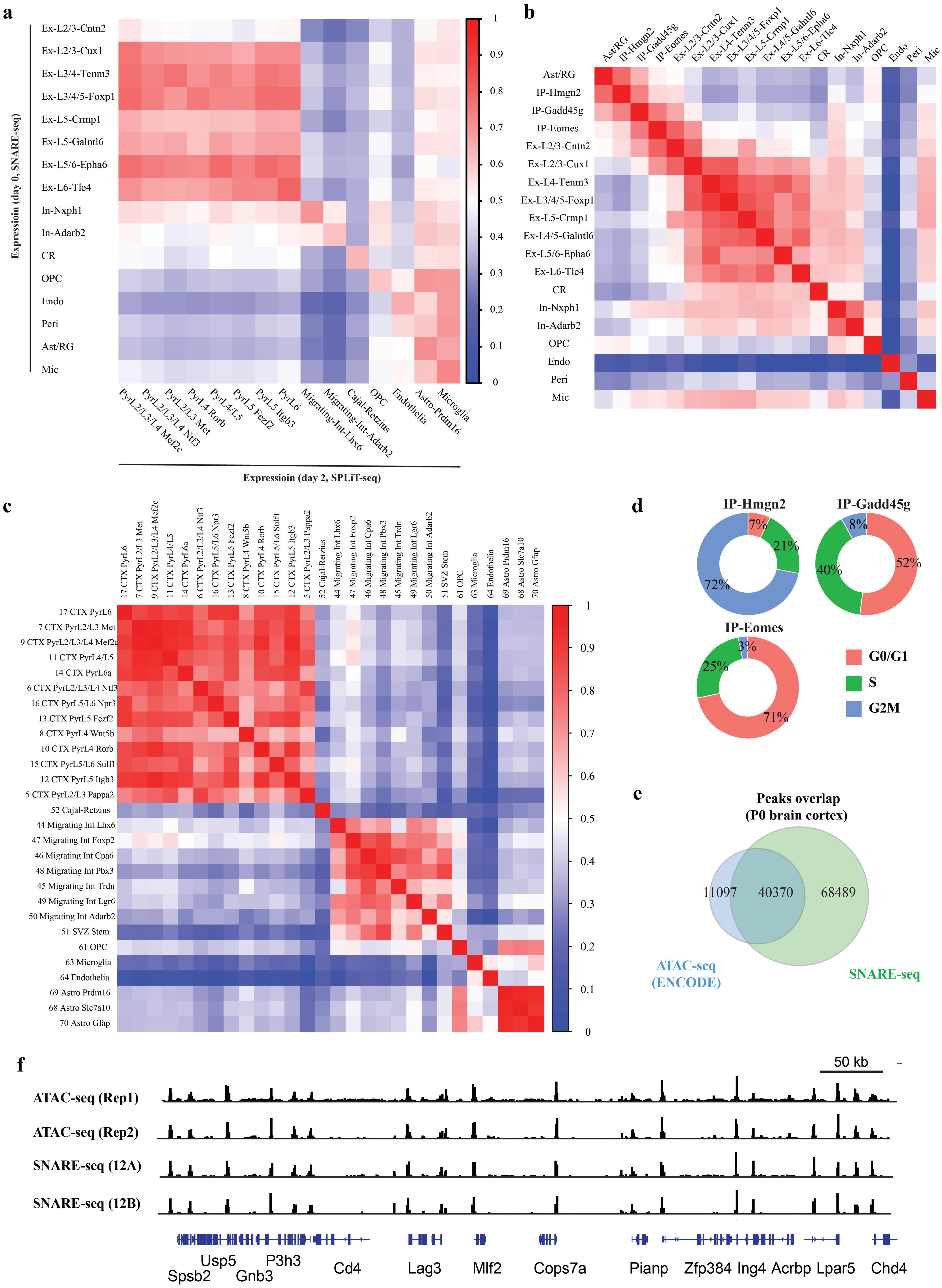

**Supplementary Figure 10.** Neonatal mouse cerebral cortex SNARE-seq profiles are correlated with published expression and chromatin data. **a**, Correlation heatmap of mouse cerebral cortex cell types identified with SNARE-seq expression data compared with previously identified cell types using SPLiT-seq. **b**, Intra-assay pair-wise correlation heatmap of cell types identified with SNARE-seq expression data. **c**, Intra-assay pair-wise correlation heatmap of cell types identified with SPLiT-seq expression data. **d**, Proportion of nuclei in different cell cycle phases showing cell cycle exit of late intermediate progenitor cells. **e**, Representative Venn diagram showing the number of overlap of common peaks between bulk ATAC-seq (ENCODE) data and SNARE-seq chromatin profiles of mouse brain cortex. **f**, Aggregated SNARE-seq chromatin profiles (bottom two tracks) agree with bulk ATAC-seq (ENCODE, top two tracks) and are consistent between independent experiments.

#### Supplementary Figure 11

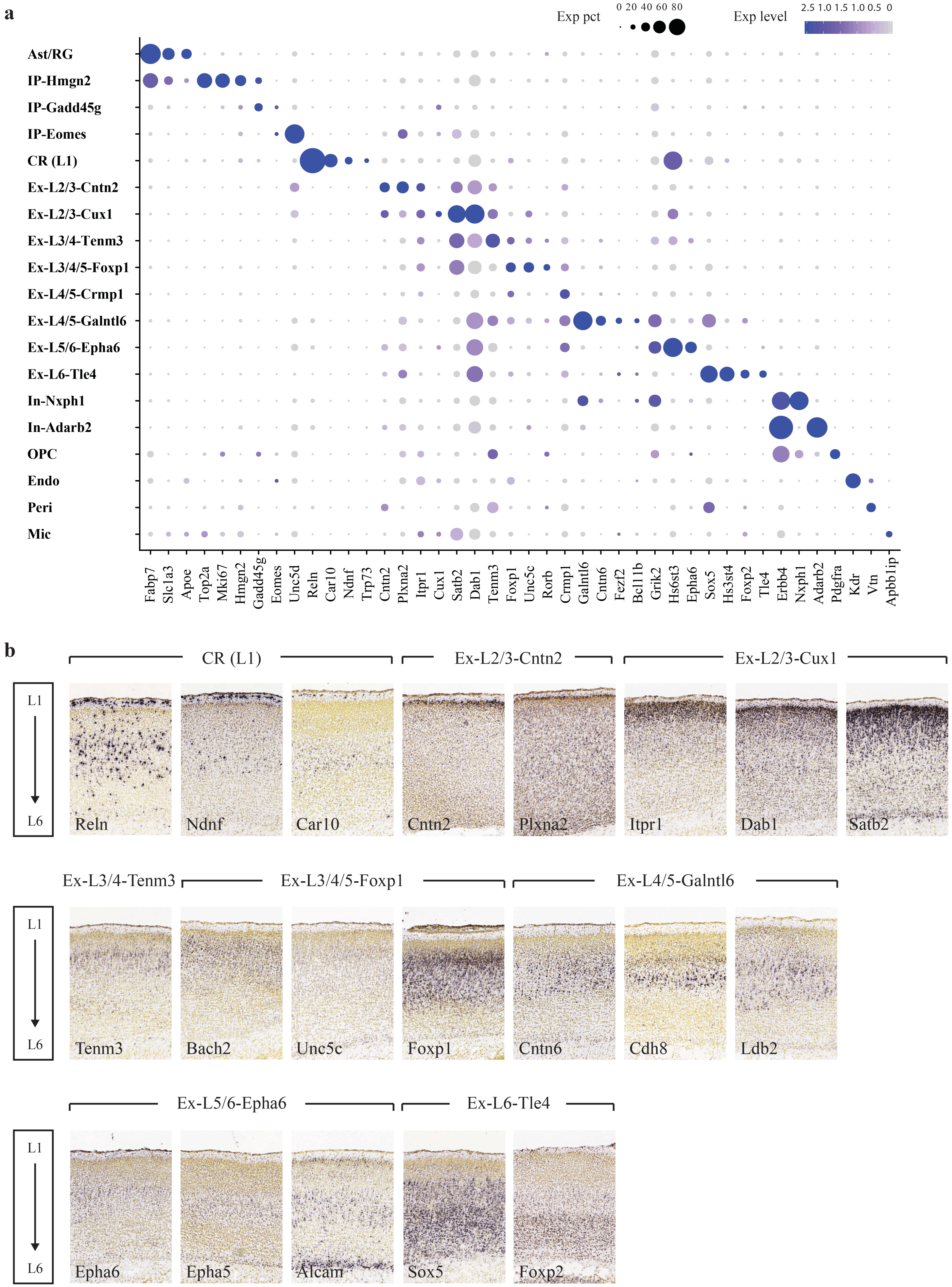

**Supplementary Figure 11.** SNARE-seq expression data identified cell type specific markers of mouse neonatal cerebral cortex. **a**, Dot plot showing the expression of known and novel marker genes (**Table S1**) in each cell type. **b**, RNA *in situ* hybridization (ISH) stains (Allen Human Brain) of postnatal (day 4) mouse cerebral cortex showing layer-specific marker gene expression.

#### Supplementary Figure 12

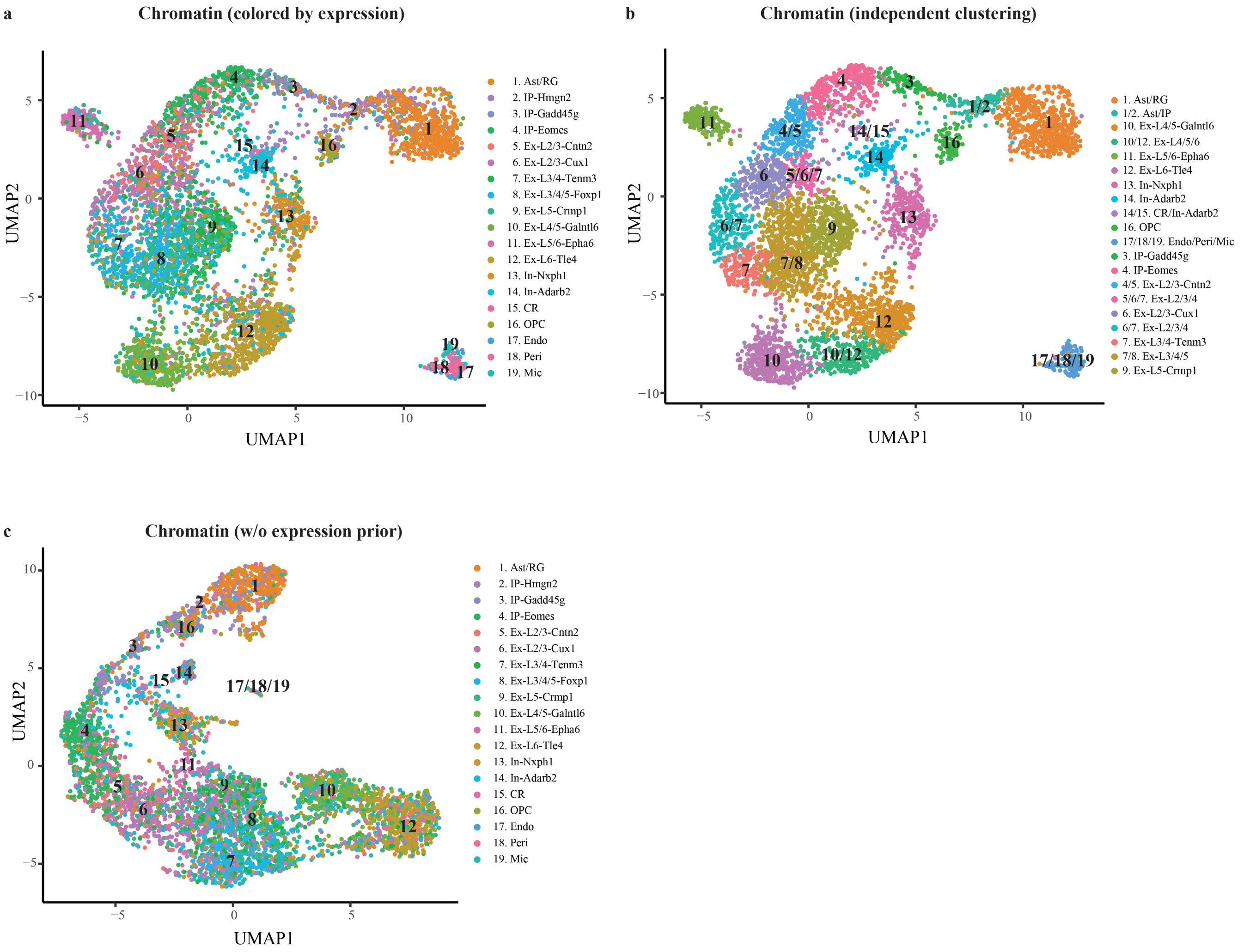


**Supplementary Figure 12.** Clustering of SNARE-seq single-cell chromatin accessibility on mouse neonatal cerebral cortex. **a**, UMAP projection of SNARE-seq chromatin data generated with aggregating chromatin reads by transcriptome labeling first, same as Figure 2c. Cells are labeled with the same color codes for cell types identified by the linked expression data. **b**, Same UMAP projection of SNARE-seq chromatin as **a**, but cells are labeled with results of independent clustering with each cell's Principal Component scores of topic information calculated by cisTopic. **c**, Different from **a** and **b**, chromatin peak count matrix were generated without using any expression information, and data was clustered independently by cisTopic. Cells are labeled with corresponding cell types identified by expression profile.

#### Supplementary Figure 13

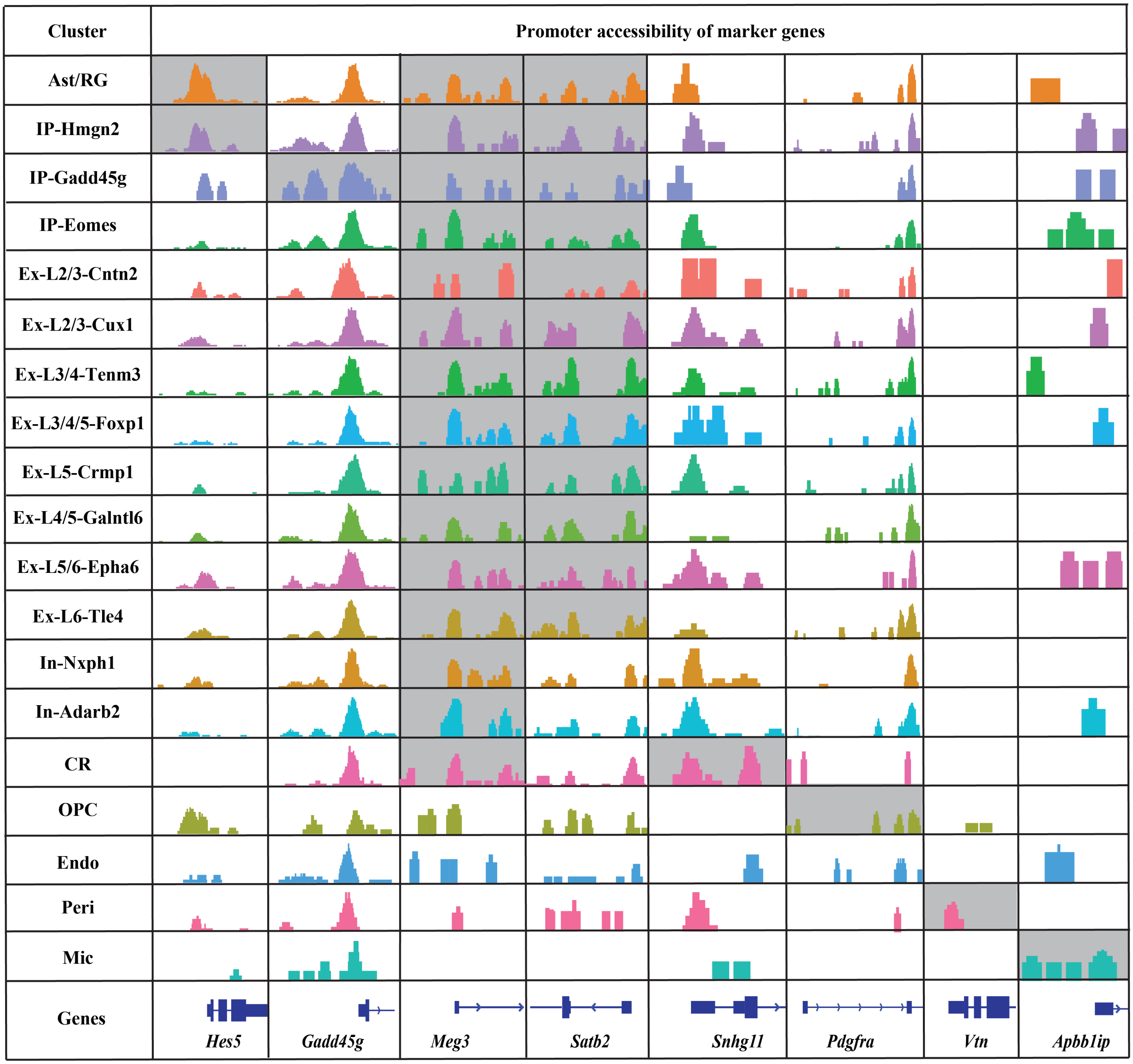


**Supplementary Figure 13.** Aggregate chromatin accessibility profiles at loci of cell type-specific marker genes. For better visualization, tracks of differential expressing genes in respective cell types are shaded in gray.

#### Supplementary Figure 14

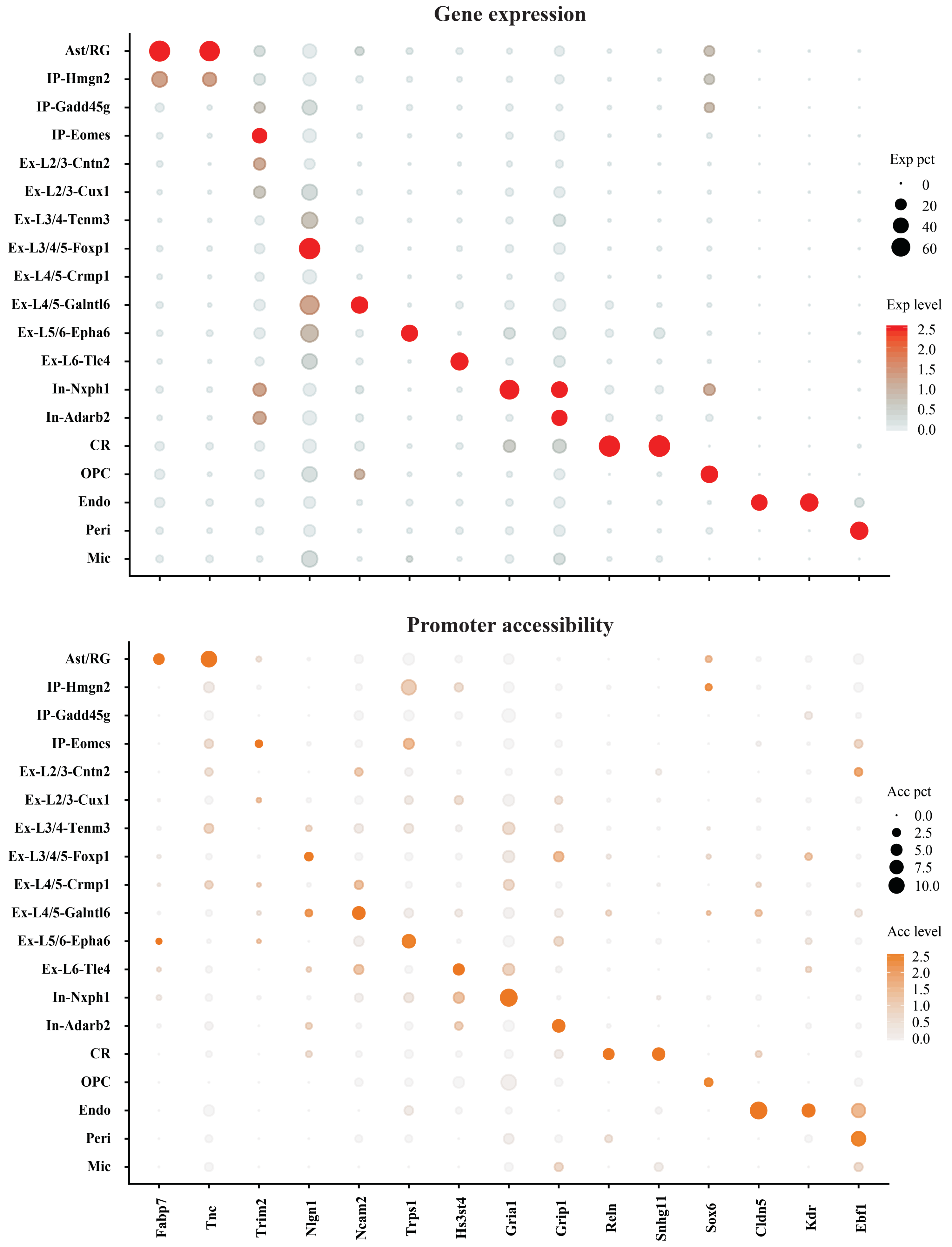

**Supplementary Figure 14.** SNARE-seq links the promoter accessibility with expression level of cell type-specific genes. Top, dot plot showing the differential expression of genes in each cell type. The size of the dot represents the percentage of nuclei within a cell type expressing the gene and the color depth indicates the average expression level. Bottom, dot plot showing the promoter accessibility of markers in each cell type. The size of the dot represents the percentage of nuclei within a cell type that is accessible in promoter regions of corresponding genes and the color depth indicates the average accessibility level.

**Supplementary Figure 15**

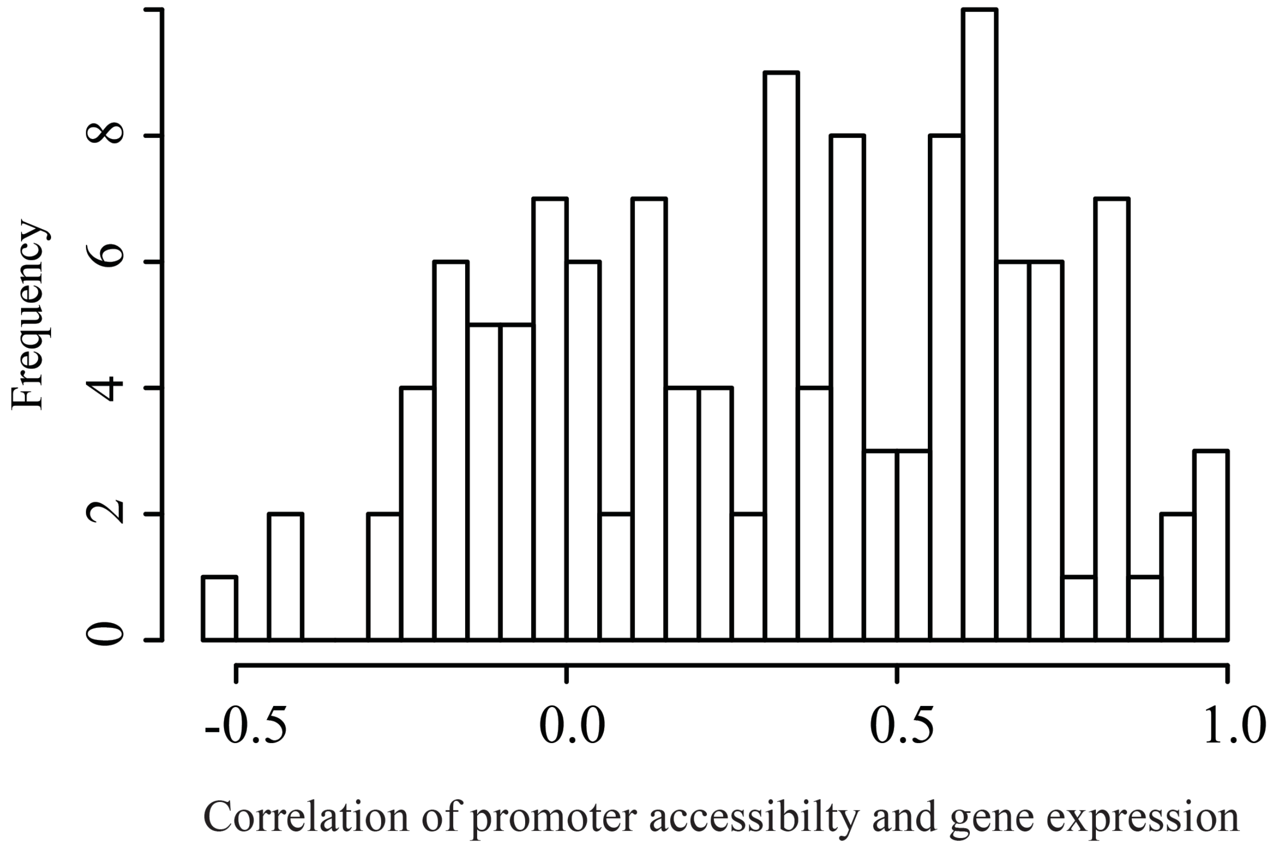


**Supplementary Figure 15**. Correlation of promoter accessibility and expression levels across all cell types for the lineage-specific genes.

**Supplementary Figure 16**

| | GO Biological Process | p-Value | Motif | TF | p-Value |
|---|---|---|---|---|---|
| **Ast/RG** | somatic stem cell maintenance | 1.41E-31 | 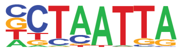 | LHX2 | 1E-142 |
| **IP-Hmgn2** | spinal cord development | 6.01E-12 | 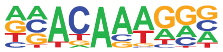 | SOX10 | 1E-60 |
| **IP-Gadd45g** | Wnt receptor signaling pathway | 2.96E-09 | 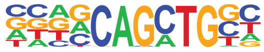 | TCF12 | 1E-45 |
| **IP-Eomes** | telencephalon development | 4.29E-12 | 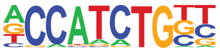 | NEUROD1 | 1E-233 |
| **Ex-L2/3-Cntn2** | ventricular cardiac muscle cell differentiation | 2.02E-06 | 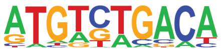 | PBX3 | 1E-14 |
| **Ex-L2/3-Cux1** | embryonic skeletal system development | 1.28E-05 | 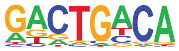 | MEIS1 | 1E-36 |
| **Ex-L3/4-Tenm3** | regulation of vasculogenesis | 2.64E-06 | 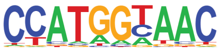 | RFX1 | 1E-53 |
| **Ex-L3/4/5-Foxp1** | response to alkaloid | 6.70E-05 | 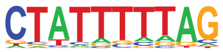 | MEF2A | 1E-34 |
| **Ex-L5-Crmp1** | protein O-linked glycosylation | 1.52E-05 | 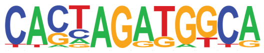 | CTCF | 1E-22 |
| **Ex-L4/5-Galntl6** | neutrophil chemotaxis | 2.17E-05 | 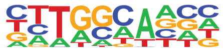 | NF1 | 1E-81 |
| **Ex-L5/6-Epha6** | actin cytoskeleton reorganization | 1.84E-05 | 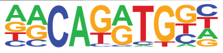 | NEUROG2 | 1E-212 |
| **Ex-L6-Tle4** | central nervous system neuron development | 3.89E-07 | 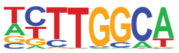 | NFIA | 1E-176 |
| **In-Nxph1** | lens development in camera-type eye | 8.76E-10 | 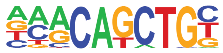 | E2A | 1E-291 |
| **In-Adarb2** | central nervous system neuron differentiation | 5.44E-07 | 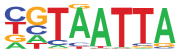 | EMX2 | 1E-49 |
| **CR** | response to light stimulus | 4.91E-07 | 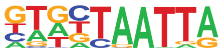 | LHX1 | 1E-49 |
| **OPC** | stem cell differentiation | 2.89E-10 | 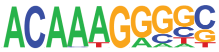 | SOX2 | 1E-54 |
| **Endo** | blood vessel morphogenesis | 5.78E-28 | 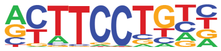 | ETV2 | 1E-171 |
| **Peri** | blood vessel morphogenesis | 6.90E-14 | 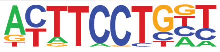 | ELF3 | 1E-50 |
| **Mic** | response to biotic stimulus | 4.58E-10 | 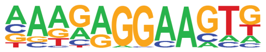 | SFPI1(PU.1) | 1E-55 |

**Supplementary Figure 16.** SNARE-seq chromatin data from mouse neonatal cerebral cortex uncovers lineage-specific regulatory information. Gene ontology analysis of top biological process using GREAT and transcription factor motifs discovered using HOMER for each cell type. Interesting examples are shaded in gray.

#### Supplementary Figure 17

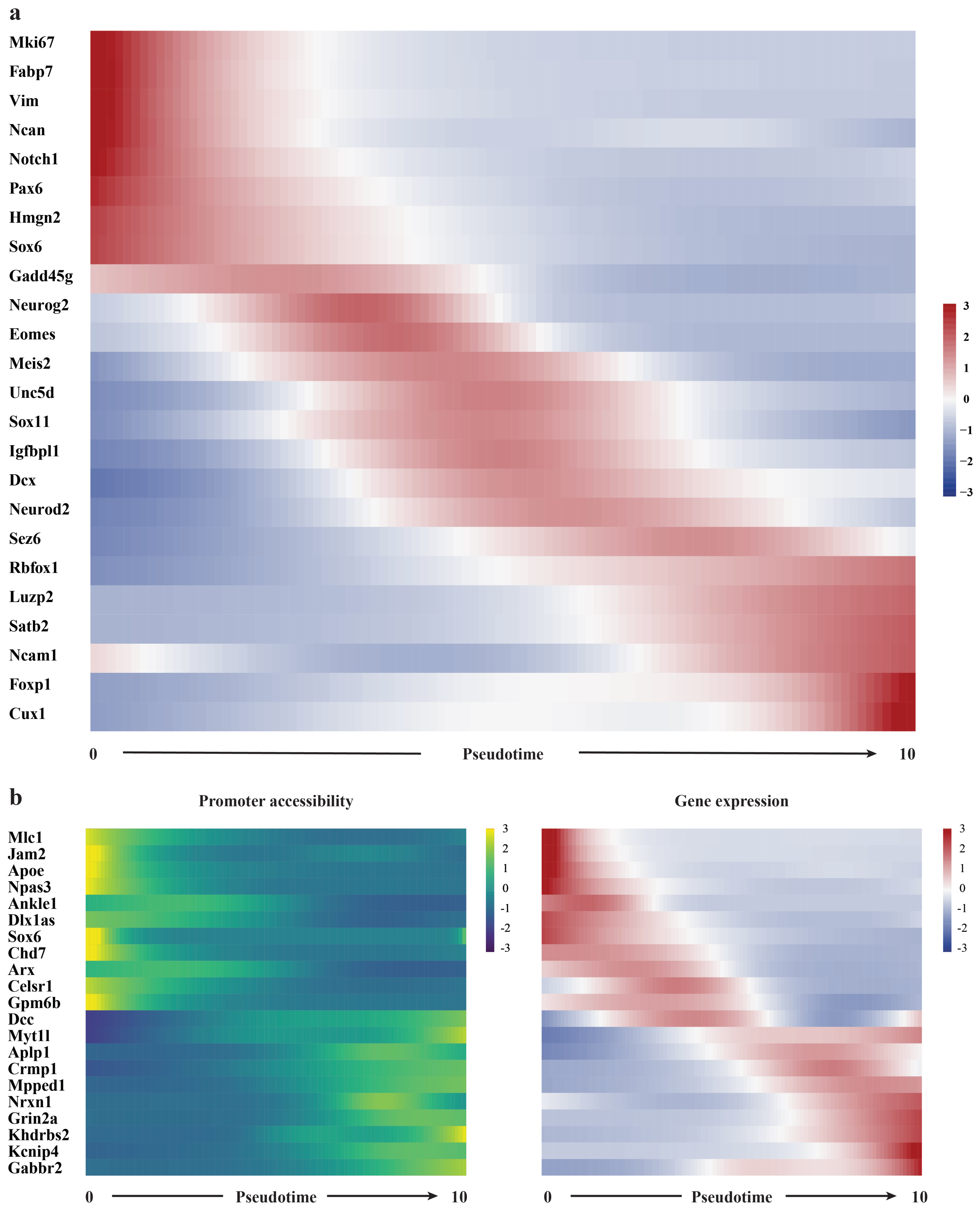


**Supplementary Figure 17.** Pseudotime analysis reveals dynamics of gene expression in early neurogenesis. **a**, Heatmap of selected genes involved in neurogenesis showing the expression changes along developmental trajectory. **b**, Promoter accessibility and gene expression dynamics showing mostly similar directional changes along pseudotime.

#### Supplementary Figure 18

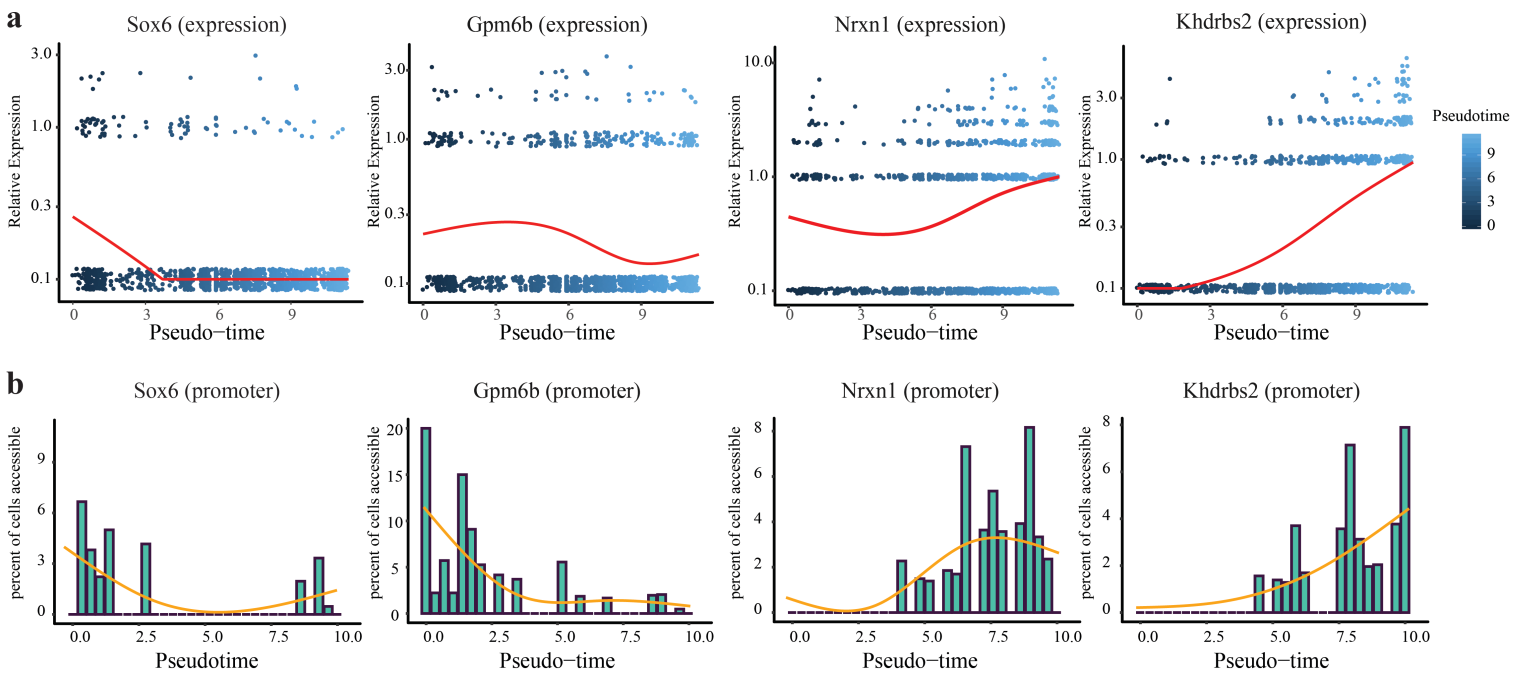


**Supplementary Figure 18. a,** Relative expression of gene markers identified by pseudotime trajectory analysis. Black line indicates the pseudotime-dependent average from a smoothed binomial regression. The relative expression is calculated by first model read count with the negative binomial and then normalize with cell size factor estimated by estimateSizeFactors function in Monocle. Smoothed pseudotime-dependent gene expression curve as shown in Figure 2g is colored in red. **b,** Histogram showing the percentage of cells that had accessible promoters calculated by aggregating chromatin signals into bins (bins=30) along pseudotime, and smoothed curve as shown in Figure 2g is colored in orange.

**Supplementary Figure 19**

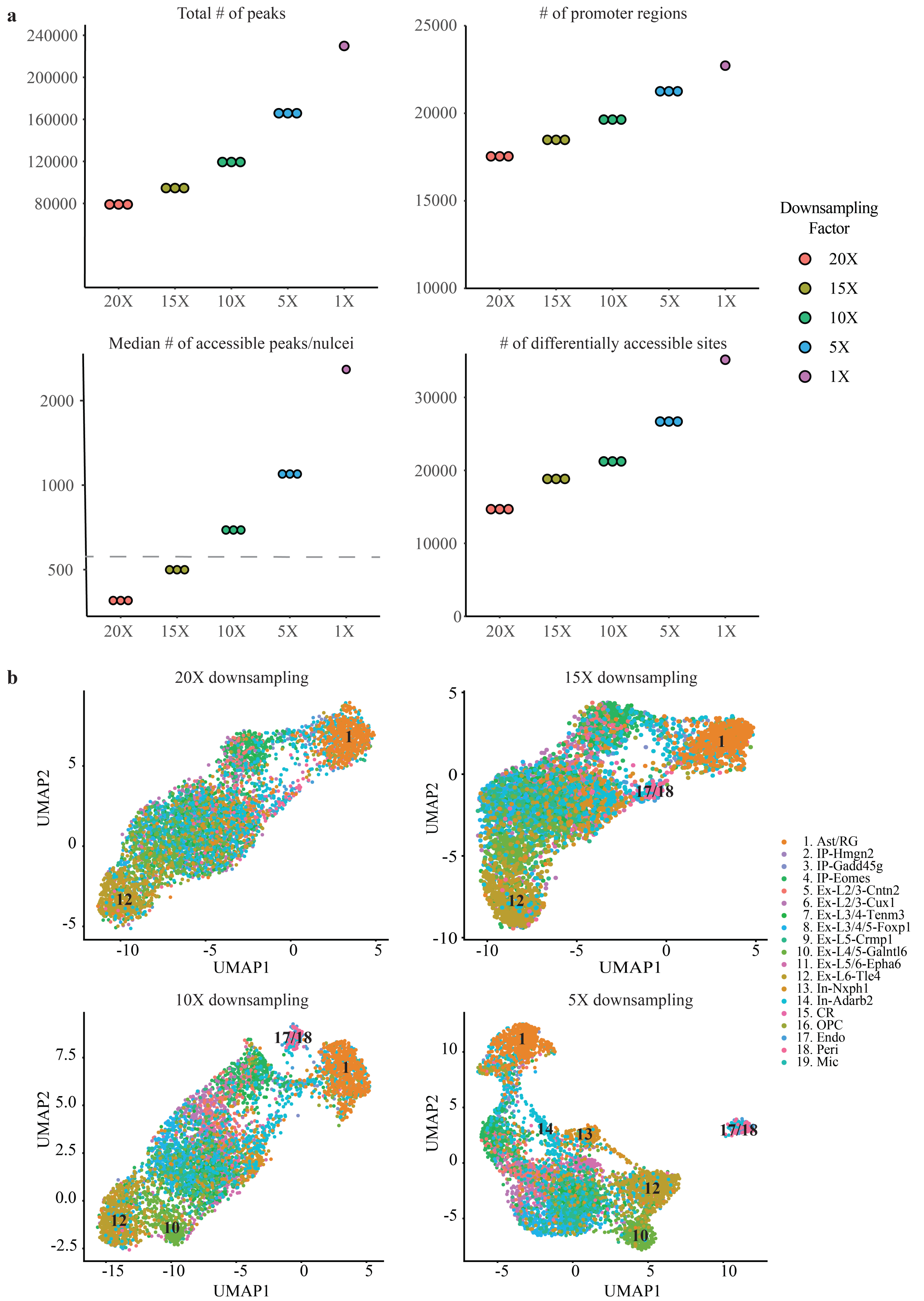

**Supplementary Figure 19.** Sensitivity of SNARE-seq chromatin data. **a,** Total called peaks, recovered promoter regions, median numbers of accessible sites detected per nuclei and total differential accessible sites across all cell types after downsampling raw reads 20-, 15-, 10- and 5-folds. Each dot represents a random downsampling test. Sci-CAR data have a median of ~700 accessible sites/nucleus (indicated by the dash line), which is close to the 15x down-sampled SNARE-seq data. **b,** Representative UMAP projection of cisTopic clustering result of SNARE-seq chromatin data of mouse neonatal cerebral cortex at varying down-sampled depths. Cells are labeled with the same color codes for cell types identified by the linked expression data and those discernible clusters are labeled with cluster identities showing on the right.

#### Supplementary Figure 20

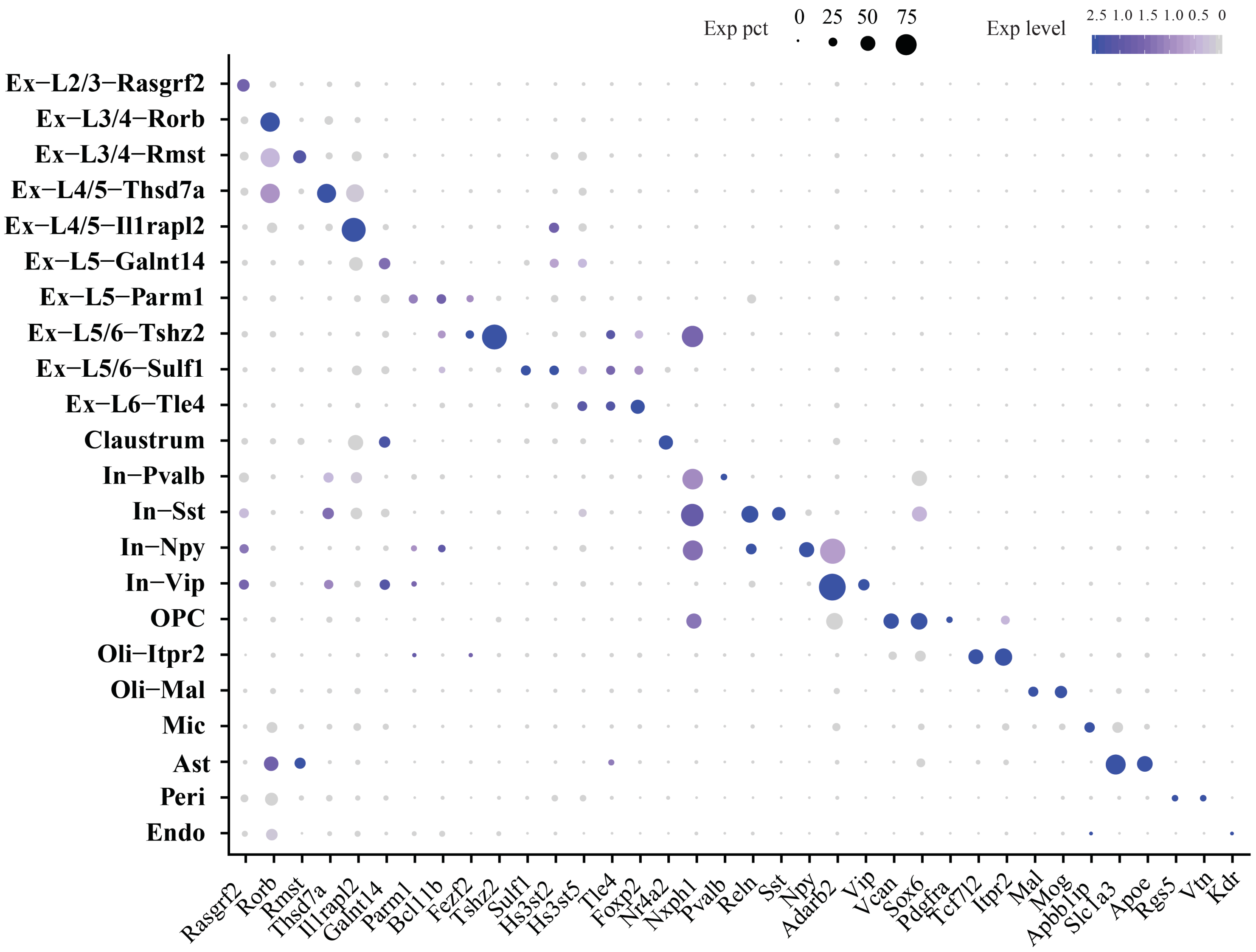


**Supplementary Figure 20.** Dot plot showing the expression of marker genes (**Table S1**) for each cell type in adult mouse cerebral cortex identified by SNARE-seq expression data.

### Supplementary Figure 21

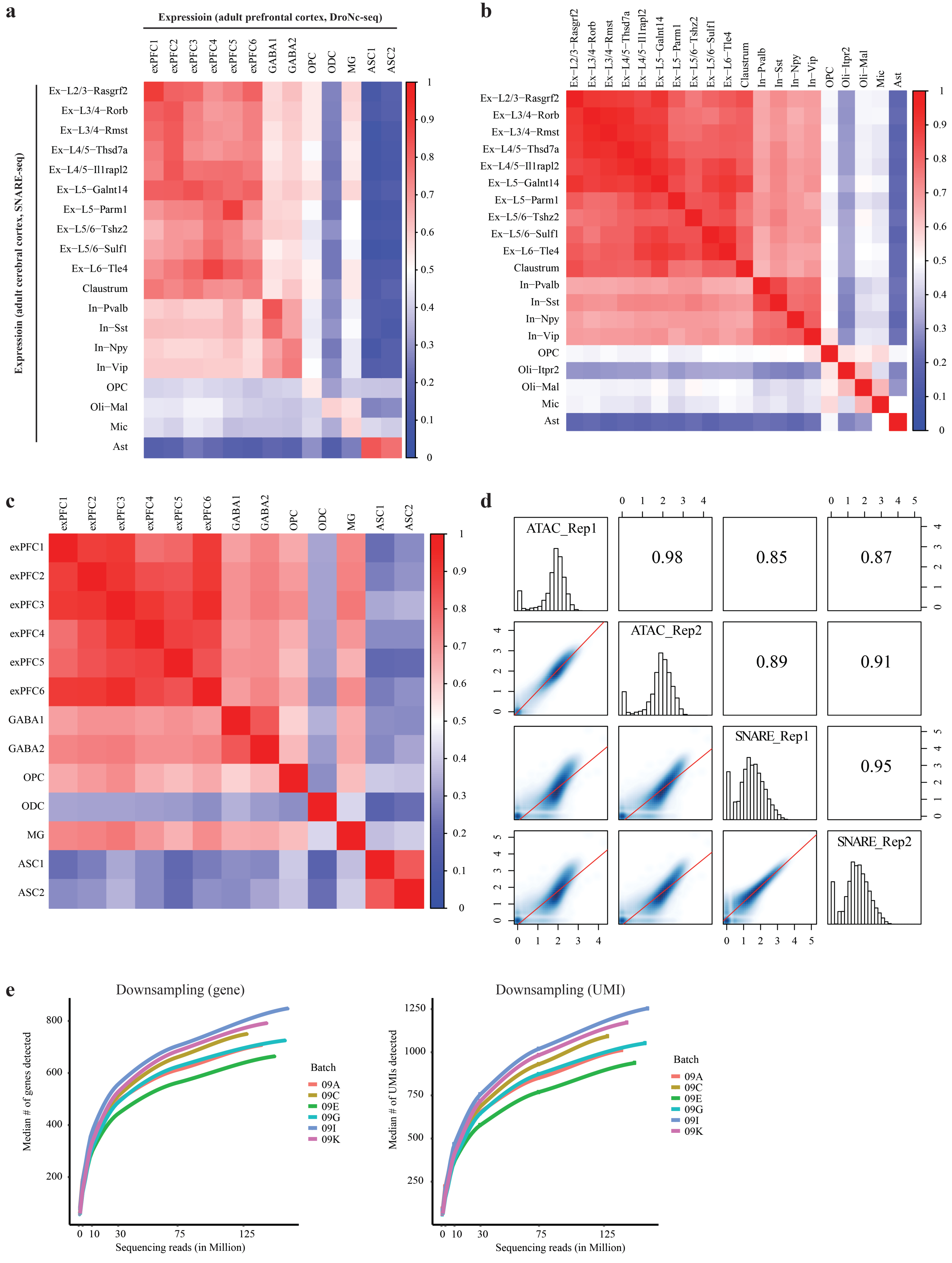

**Supplementary Figure 21.** The SNARE-seq profiles of adult mouse cerebral cortex are correlated with published expression and chromatin data. **a**, Correlation heatmap of mouse cerebral cortex cell types identified with SNARE-seq expression data compared with previously identified cell types using DroNc-seq. **b**, Intra-assay pair-wise correlation heatmap of cell types identified with SNARE-seq expression data. **c**, Intra-assay pair-wise correlation heatmap of cell types identified with DroNc-seq expression data. **d**, Pair-wise correlation of chromatin accessibility profiles between adult mouse frontal cortex replicates (ENCODE) and SNARE-seq replicates. Aggregated genome coverage was log10 normalized. **e**, In silico downsampling showing the change of genes and UMIs detected by SNARE-seq expression on different sequencing depth.

**Supplementary Figure 22**

| | GO Biological Process | p-Value | Motif | TF | p-Value |
|---|---|---|---|---|---|
| **Ex-L2/3-Rasgrf2** | inactivation of MAPK activity | 4.9E-14 | 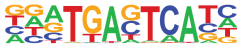 | FRA1 | 1E-286 |
| **Ex-L3/4-Rorb** | unsaturated fatty acid metabolic process | 4.6E-09 | 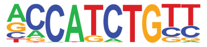 | NEUROG2 | 1E-233 |
| **Ex-L3/4-Rmst** | regulation of neurological system process | 2.3E-06 | 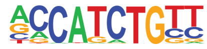 | NEUROG2 | 1E-65 |
| **Ex-L4/5-Thsd7a** | regulation of peptide hormone secretion | 3.5E-09 | 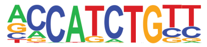 | NEUROG2 | 1E-360 |
| **Ex-L4/5-Il1rapl2** | heparan sulfate proteoglycan metabolic process | 8.9E-06 | 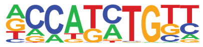 | OLIG2 | 1E-125 |
| **Ex-L5-Galnt14** | cartilage development | 1.3E-06 | 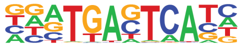 | FRA1 | 1E-110 |
| **Ex-L5-Parm1** | smooth muscle contraction | 6.7E-07 | 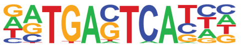 | ATF3 | 1E-360 |
| **Ex-L5/6-Tshz2** | organelle localization | 7.2E-09 | 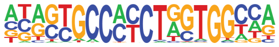 | CTCF | 1E-76 |
| **Ex-L5/6-Sulf1** | neurotransmitter transport | 3.8E-05 | 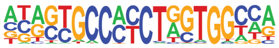 | CTCF | 1E-28 |
| **Ex-L6-Tle4** | response to alkaloid | 2.7E-14 | 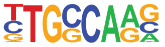 | NF1 | 1E-235 |
| **Claustrum** | exocytosis | 7.5E-09 | 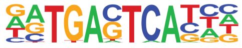 | ATF3 | 1E-167 |
| **In-Pvalb** | actin filament bundle assembly | 1.9E-10 | 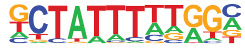 | MEF2B | 1E-260 |
| **In-Sst** | synapse organization | 8.0E-09 | 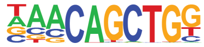 | TCF21 | 1E-187 |
| **In-Npy** | neuromuscular process | 2.7E-10 | 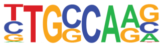 | NF1 | 1E-39 |
| **In-Vip** | calcium ion transmembrane transport | 1.7E-07 | 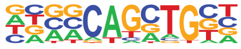 | ASCL1 | 1E-46 |
| **OPC** | small GTPase mediated signal transduction | 2.0E-08 | 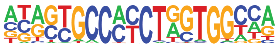 | CTCF | 1E-30 |
| **Oli-Itpr2** | actin filament-based process | 4.5E-07 | 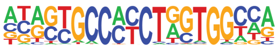 | CTCF | 1E-53 |
| **Oli-Mal** | axon ensheathment | 2.1E-14 | 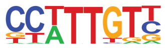 | SOX3 | 1E-167 |
| **Mic** | endocytosis | 1.2E-09 | 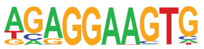 | PU.1 | 1E-31 |
| **Ast** | regulation of Rho protein signal transduction | 5.3E-18 | 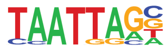 | LHX2 | 1E-188 |
| **Peri** | regulation of exocytosis | 4.4E-05 | 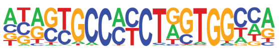 | CTCF | 1E-34 |
| **Endo** | neuromuscular process controlling balance | 1.8E-06 | 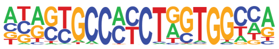 | CTCF | 1E-58 |

**Supplementary Figure 22.** Gene ontology of top biological process and transcription factor motifs analysis of each cell type in adult mouse cerebral cortex using SNARE-seq chromatin data.
